# A Novel ACLY Inhibitor Induces Transcriptional and Metabolic Reprogramming to Overcome Tyrosine kinase inhibitor Resistance in Lymphoma

**DOI:** 10.64898/2026.09.29.755331

**Authors:** Nagarajan Pattabiraman, Cosimo Lobello, David Rushmore, Shengchun Wang, Luca Mologni, Hossein Borghaei, Kavindra Nath, Wujuan Zhang, Aaron R. Goldman, Reza Nejati, Mariusz Wasik, Johnvesly Basappa

**Affiliations:** Veracure Biosciences Inc., Gaithersburg, MD; Fox Chase Cancer Center, Philadelphia, PA, USA; Department of Medicine and Surgery, University of Milano-Bicocca, Monza, Italy; Proteomics & Metabolomics Shared Resource, The Wistar Institute, Philadelphia, PA, USA; Department of Radiology, University of Pennsylvania, Philadelphia, PA, USA

**Keywords:** ACLY, tyrosine phosphorylation, lipid metabolism, drug discovery

## Abstract

**Background:** Metabolic plasticity contributes to therapeutic resistance in T-cell lymphoma, but the metabolic dependencies underlying tyrosine kinase inhibitor (TKI) resistance remain poorly understood. ATP citrate lyase (ACLY), which converts citrate to cytosolic acetyl-CoA, links glucose metabolism to lipogenesis and acetylation-dependent regulation. We hypothesized that ACLY represents an actionable metabolic vulnerability in lymphoma.

**Methods:** Structure-guided drug discovery and biochemical screening were used to identify the novel ACLY inhibitor VCB1. VCB1 was evaluated against recombinant human ACLY and in TKI-sensitive and TKI-resistant lymphoma models. Antitumor activity was assessed by proliferation, cell-cycle analysis, apoptosis, and immunoblotting, followed by integrated transcriptomic, metabolomic, and lipidomic profiling.

**Results:** Our xenograft study showed that ACLY dependence in ALK+ALCL. We discovered novel ACLY inhibitor VCB1 directly inhibited recombinant ACLY with µM potency (IC₅₀, 0.9 µM) and strongly suppressed lymphoma cell growth. In ALK inhibitor-sensitive SUPM2, VCB1 showed an IC₅₀ of 0.299 μM and retained potency in lorlatinib-resistant SUPM2-LR1000 cells (1.62 μM), whereas BMS-303141 was substantially less active (17.46 and 22.07 μM). VCB1 similarly inhibited BTK inhibitor sensitive and resistant mantle cell lymphoma cell growth (IC₅₀, 0.28 and 0.278 μM), compared with BMS-303141 (30.5 and 37.01 μM). Multi-omics profiling revealed suppression of de novo lipogenesis, including *ACACA, FASN*, and *PPARGC1B*, together with extensive remodeling of triglyceride, phospholipid, cardiolipin, ceramide, and acylcarnitine species and increased expression of fatty-acid oxidation genes. VCB1 activated the integrated stress response (ISR), inducing *ATF4, PPP1R15A, DDIT3, DDIT4* and *SESN2*, while disrupting purine and pyrimidine metabolism and reducing nucleotide biosynthetic genes, including *CAD, DHODH, UCK2*, and *ATIC*. VCB1 further activated an ACLY–ISR–CHAC1 axis associated with reduced GCLC/GCLM expression, glutathione depletion, and a decreased GSH/GSSG ratio, indicating impaired redox homeostasis. TKI-resistant cells exhibited distinct lipid-metabolic remodeling consistent with metabolic plasticity. VCB1 also altered immune-regulatory programs involving *PDCD1, TIGIT, CD247, CD226, NFATC2*, and inflammatory cytokines. Functionally, VCB1 induced cell cycle arrest and apoptosis.

**Conclusions:** ACLY represents an integrated metabolic and signaling vulnerability in lymphoma. VCB1 suppresses lymphoma growth across TKI-sensitive and resistant models by disrupting lipid and nucleotide metabolism, metabolic adaptation and ISR and redox homeostasis supporting ACLY inhibition as a therapeutic strategy for TKI-resistant lymphoma.

## Background

The global rise in cardiometabolic disorders, including atherosclerosis, non-alcoholic fatty liver disease (NAFLD), and type 2 diabetes, has intensified efforts to identify therapeutic targets beyond conventional lipid-lowering strategies. ATP-citrate lyase (ACLY) is a cytosolic enzyme that occupies a central position in cellular metabolism by catalyzing the ATP-dependent conversion of citrate into acetyl-CoA and oxaloacetate. As the primary source of cytosolic acetyl-CoA, ACLY supports de novo lipid biosynthesis and histone acetylation, thereby integrating carbohydrate metabolism with lipid synthesis and epigenetic regulation(1–3). ACLY activity is tightly regulated by multiple post-translational modifications, including phosphorylation, acetylation, ubiquitination, and glycosylation. Among these, phosphorylation of serine 455 enhances enzymatic activity (4). We previously identified tyrosine 682 phosphorylation of ACLY as a novel regulatory event mediated by oncogenic tyrosine kinases and demonstrated its functional impact on ACLY activity (5, 6). These modifications increase ACLY-dependent acetyl-CoA production, thereby promoting lipid biosynthesis and metabolic reprogramming. Consistently, ACLY overexpression and hyperactivation have been implicated in diverse pathological conditions, including hyperlipidemia (7), non-alcoholic fatty liver disease (NAFLD)(8, 9), and multiple cancers, including lung(10), breast(11), prostate(12), liver(13), pancreatic(14) and colon cancer(15), where enhanced acetyl-CoA metabolism supports tumor growth, survival, and progression.

Emerging evidence has established ACLY as a metabolic vulnerability across cancer types. ACLY-derived acetyl-CoA directly fuels histone acetylation, thereby linking cellular metabolism to chromatin remodeling and epigenetic plasticity. This is particularly relevant in hematologic malignancies, which are characterized by profound metabolic and epigenetic rewiring. In murine hematopoietic stem and progenitor cells (HSPCs), pharmacologic inhibition or genetic deletion of ACLY promotes myeloid differentiation (16). In PTEN-deficient T-cell acute lymphoblastic leukemia (T-ALL), ACLY is required for thymocyte transformation and tumor growth (17), while cooperative interactions between ACLY and MYC further drive leukemogenesis (18). In acute myeloid leukemia (AML), ACLY overexpression accelerates disease progression (19). Moreover, genetic or pharmacologic inhibition of ACLY suppresses tumor growth, induces apoptosis, and enhances therapeutic sensitivity in leukemia and lymphoma models (20). Collectively, these findings position ACLY as both a metabolic regulator and epigenetic modulator with therapeutic relevance in hematologic malignancies.

Despite decades of research, the development of clinically effective ACLY inhibitors has proven challenging. Early-generation inhibitors, including SB-204990 (21, 22), BMS-303141(23), and the recently reported NDI-091143(24), demonstrated potent enzyme inhibition but were limited by poor aqueous solubility, suboptimal oral bioavailability, short in vivo half-lives, and insufficient cellular activity. These pharmacologic shortcomings restricted their translation into effective systemic therapies. Although ACLY has attracted considerable interest as a therapeutic target for hyperlipidemia, cancer, and other metabolic disorders, the discovery of clinically viable inhibitors has been hindered by the predominance of substrate- like compounds targeting conserved catalytic pockets and the limited availability of chemically diverse scaffolds with favorable pharmacological properties(25). The clinical approval of Bempedoic acid by the U.S. Food and Drug Administration provided the first validation of ACLY as a druggable metabolic target in humans as a cholesterol lowering agent (26). Bempedoic acid is a liver-specific pro-drug activated by ACSVL1, an enzyme predominantly expressed in hepatocytes, where it inhibits ACLY and reduces acetyl-CoA production required for cholesterol and lipid biosynthesis (27). More recently, EVT0185 was identified as an ACLY inhibitor that is similarly converted to its active CoA derivative by the liver-enriched enzymes FATP2 and ACSVL1(28). While these compounds demonstrate the therapeutic feasibility of ACLY inhibition, their liver-restricted activation limits broader applications, particularly in oncology, where effective systemic inhibition is necessary to target malignant cells residing in diverse tissues. Despite these advances, most ACLY inhibitors reported to date exhibit limitations related to pharmacokinetic properties, tissue-selective activation, or insufficient cellular activity, underscoring the need for new compounds capable of achieving effective systemic ACLY inhibition in cancer. Such efforts are particularly relevant in hematologic malignancies, where ACLY-mediated metabolic and epigenetic programs contribute to disease progression and therapeutic resistance.

Building upon the clinical validation of ACLY inhibition and recent advances in high-resolution ACLY structural biology (29), we employed a structure-based virtual screening strategy to identify novel small-molecule ACLY inhibitors with improved cellular activity. Given the dual role of ACLY in regulating metabolic pathways and acetylation-dependent gene expression, we investigated the biological consequences of ACLY inhibition in ALK-positive anaplastic large cell lymphoma (ALK+ ALCL) models, including both ALK inhibitor sensitive (ALKi-S) and resistant (ALKi-R) cell lines. This work builds upon our earlier discovery that oncogenic NPM-ALK directly phosphorylates ACLY at tyrosine 682, thereby regulating lipid metabolism and promoting ALCL tumorigenesis(6, 30, 31). Subsequent studies established that ACLY Y682 is also a target of Src-family kinases, including LYN, and identified ACLY as a novel signaling effector downstream of the PI3K/PIP2/PIP3/LYN axis in acute myeloid leukemia, linking oncogenic kinase signaling to acetyl-CoA production, lipid biosynthesis, histone acetylation, and leukemic cell growth(32). These findings, together with translational efforts targeting the PI3K/LYN/ACLY signaling network (WO 2020/243063 A1), provided the biological rationale for the development of direct ACLY inhibitors described herein. Our studies provide mechanistic insights into ACLY dependent metabolic regulation and establish a framework for the development of next-generation ACLY targeted therapies for hematologic malignancies.

In this study, we build upon the signaling–metabolism axis by identifying and characterizing novel ACLY inhibitors through structure-based virtual screening and evaluating their activity in ALK+ ALCL models with distinct ALKi sensitivity and resistance. Using integrated transcriptomic, metabolomic, and untargeted lipidomic profiling, we define the metabolic and transcriptional consequences of ACLY inhibition and identify stress-response pathways associated with therapeutic response. These findings establish ACLY as an actionable metabolic vulnerability and provide a mechanistic framework for developing next-generation ACLY-targeted strategies for TKI-resistant hematologic malignancies.

## Methods and materials

### Cell lines

### ALK Inhibitor (ALKi)-Resistant ALCL Cell Lines

ALCL cell lines cultured as described by our group earlier(33, 34). Briefly, SUPM2 parental and ALK inhibitors crizotinib-and lorlatinib resistant SUPM2 cell lines SUPM2 CR03 (resistant up to 300 nM of crizotinib and cultured at that concentration) and ALKi lorlatinib-resistant cell line SUPM2 LR1000 (resistant up to 1000 nM of lorlatinib and cultured at 300 nM of lorlatinib). The cell lines were grown in RPMI medium supplemented with 10% FBS and 1% penicillin/streptomycin in a humidified incubator at 37 °C with 5% CO_2_. For the MTT assay the cell lines were culture in were grown in RPMI medium supplemented with 5% FBS and 1% penicillin/streptomycin in a humidified incubator at 37 °C with 5% CO_2_Mantle cell lymphoma (MCL) cell lines JeKo-1 and Maver were cultured in RPMI-1640 medium supplemented with 10% fetal bovine serum (FBS) and 1% penicillin/streptomycin (Pen/Strep) in a humidified incubator at 37 °C with 5% CO_2_. For the MTT assay the cell lines were culture in were grown in RPMI medium supplemented with 5% FBS and 1% penicillin/streptomycin in a humidified incubator at 37 °C with 5% CO_2._

### Reagents and antibodies

Cell Proliferation Reagent WST-1 (Millepore-sigma), CellTiter 96® Non-Radioactive Cell Proliferation Assay (MTT) kit (catalog#G4001) from Promega. HA Pierce IP Lysis Buffer, Halt Protease and Phosphatase Inhibitor, GAPDH (catalog# 60004-1-Ig) from Proteintech, HA-Tag (C29F4) Rabbit (catlog# 3724S), Phospho-ALK (Tyr1604) Antibody (catalog# 3341S), ALK (D5F3®) XP® Rabbit (catalog# 3633S) from Cell Signaling Technology. ACLY antibody from Proteintech.

### In-silico structure based ACLY inhibitors screening

A five-point 3D pharmacophore model was generated from the adenine nucleoside interactions of ADP bound to human ATP-citrate lyase (ACLY). The model consisted of one hydrogen-bond donor (Don), three hydrogen-bond acceptors (Acc), and one aromatic ring center (Ar). Distance constraints between donor and acceptor features were incorporated to capture the essential binding interactions within the human ACLY– ADP complex. A five-point 3D pharmacophore model was generated based on the adenine nucleoside interactions observed between bound ADP and human ACLY (PDB ID: 3PFF; https//www.rcsb.org). The pharmacophore incorporated the following features: hydrogen bond donor (Don), hydrogen bond acceptor (Acc), aromatic ring center (Ar), and donor/acceptor (Don or Acc) functionalities. Distance constraints were applied between the acceptor, donor, and donor/acceptor features to preserve the spatial arrangement critical for ligand recognition. The NCI database was downloaded in its entirety and filtered prior to screening using the following criteria: molecular weight (MW) < 500 Da, fewer than 8 rotatable bonds, fewer than 5 hydrogen bond donors, and fewer than 10 hydrogen bond acceptors. Only compounds represented as a single molecular fragment were retained. Conformational analysis of the filtered compounds was subsequently carried out using the conformational search module in Molecular Operating Environment (MOE) by systematically varying all rotatable single bonds. A conformer database containing only energetically favorable conformations was then generated.

Ligand-based silico screening was performed using selected pharmacophore queries derived from the 3D pharmacophore model. Distance constraints between donor, acceptor, and donor/acceptor features were incorporated into the search criteia. The energetically favorable conformers in the NCI database were screened to identify compounds exhibiting greater than 65% feature overlap with the pharmacophore model.

### RNA extraction, library preparation and sequencing

Total RNA was isolated using the RNeasy Mini Kit (QIAGEN, Cat. no. 74104) according to the manufacturer’s instructions. RNA quantity and integrity were assessed by NanoDrop and Agilent Bioanalyzer and 400 ng of total RNA per sample was submitted to Novogene (Beijing, China) for library preparation and sequencing.

mRNA was purified from total RNA using poly-T oligo-attached magnetic beads. Following fragmentation, first-strand cDNA was synthesized using random hexamer primers, followed by second-strand cDNA synthesis. Libraries were completed through end repair, A-tailing, adapter ligation, size selection, amplification and purification, and were quantified by Qubit and real-time PCR, with size distribution assessed on a bioanalyzer. Libraries were pooled according to their effective concentration and target data output and sequenced on an Illumina platform in paired-end mode.

### Reverse transcription quantitative polymerase chain reaction

SUPM2 parental and SUPM2-LR1000 lorlatinib cell lines, were seeded in six-well plates up to 48 h. RNA isolation and DNase treatment were performed according to the kit protocol (Qiagen #74104, #79254). The RNA concentration was measured, and 1 μg was used for cDNA synthesis, which was performed with SuperScript IV (Invitrogen #18091050). For the quantitative reverse transcription polymerase chain reaction (qRT-PCR), 1 μL of cDNA was utilized. Each reaction contained: 1 μL of cDNA, 5 μL of PowerUp™ SYBR™ Green Master Mix (Applied Biosystem, A25742), 0.5 μL of forward primer (from a 10 mM stock), 0.5 μL of reverse primer (from a 10 mM stock), and 3 μL of dH2O. Each condition was tested in technical triplicate. Primers used are as follows.

Primer sequence 5′ to 3’

*ACSS1*, Forward-TTG AGA GCA CCC CAG TTT ATC; Reverse-GCA TCA CCG TAT TTC AGC AAC.

*ACSS2*, Forward-GGA TCA CTG GTC ATT CCT AC; Reverse-GTG CTG TGT AGA ACT TGG TC.

*CAD*, Forward-AGT GGT GTT TCA AAC CGG CAT; Reverse-CAG AGG ATA GGT GAG CAC TAA

*DHODH*, Forward-CCACGGGAGATGAGCGTTTC; Reverse-CAGGGAGGTGAAGCGAACA.

*PPP1R15A/GADDA34*,Forward-ATGATGGCATGTATGGTGAGC; Reverse-AACCTTGCAGTGTCCTTATCAG

*NLRP3*, Forward-GATCTTCGCTGCGATCAACAG; Reverse-CGTGCATTATCTGAACCCCAC

### Kyoto Encyclopedia of Genes and Genomes Pathway Analysis (Gene Ontology and Pathway Analysis)

The RNA sequencing (RNA-seq) data for SUPM2 and SUPM2-LR1000 ALKi sensitive and resistant cell lines were filtered for differentially expressed genes (DEG) (fold change >1; FDR < 0.05) and these datasets were subjected to Kyoto Encyclopedia of Genes and Genomes (KEGG) pathway analysis using DAVID platform.

### Bioinformatics analysis

Raw reads in FASTQ format were processed with fastp to remove adapter-containing reads, poly-N reads and low-quality reads, and Q20, Q30 and GC content of the resulting clean reads were calculated. All downstream analyses were performed on high-quality clean reads. Genome hg38 was used as reference genome. An index of the reference genome was built with HISAT2 (v2.2.1), which was used to align the paired-end clean reads with splice-aware mapping. Read counts per gene were obtained with feature Counts (v2.0.6), and gene expression levels were expressed as FPKM. Differential expression analysis between two groups was performed using DESeq2 (v1.42.0), with adjusted p-values by the Benjamini–Hochberg method; genes with an adjusted P ≤ 0.05 and log2 (fold change) ≥ 1 were considered significantly differentially expressed.

### ACLY activity assay

ACLY enzyme activity was determined using the malate dehydrogenase (MDH)-coupled method as described earlier with little modification(6, 30, 32). For the screening of potential ACLY inhibitors we used purified human recombinant ACLY protein that was incubated with various concentrations of our novel compounds (VCB1, VCB2 and VCB7 along with reference compound BMS). For ACLY activity, 500 ng to 1 µg of recombinant human proteins for each condition in triplicate well (96 plate) were incubated in reaction buffer containing 10 mM potassium citrate, 10 mM MgCl2, 1 mM DTT, 10 U malic dehydrogenase, 0.3 mM CoASH, 0.1 mM NADH in 50 mMTris (pH 8.0) and the reaction was initiated by adding 0.2mM ATP in a final volume of 100 ul, incubated at 37°C, and NADH oxidation was continuously monitored every 2 min for 60 min using a BioTeK plate reader. To measure the effect of a small molecule inhibitor of VCB1 and BMS on ALK+ALCL and MCL cell lines cells were pretreated with VCB1 and BMS at 1 µM for 24 hours. For the control experiments, lysis buffer was used in place of cell lysates for nonspecific NADH oxidation. The relative ACLY activities were calculated by normalization to the total protein abundance of the extracts in triplicate. The relative activity was calculated using PRISM GraphPad 10.6.1.

### Immunoblotting analysis

Proteins were extracted using Thermo Scientific™ Pierce™ IP Lysis Buffer (Catalog No. PI87788) containing a cocktail of protease and phosphatase inhibitors. For western blotting, 50 μg of total cell proteins were subjected to SDS-PAGE in 10 or 4-20 % NuPAGE gradient gels under reducing conditions and transferred onto a nitrocellulose membrane. The blots were blocked in 5 % skimmed milk in TBST and probed with primary antibodies overnight. The blots were developed using ECL western blotting detection reagent (GE Healthcare).

### MTT enzymatic conversion assay

Cells were plated in 96-well plates (3–5×10^3^/well) and treated with ACLY inhibitors VCB1, VCB2, VCB7 and BMS or the drug vehicle and labeled with MTT (Promega). Well contents were solubilized and absorbance was determined using a BioTeK plate reader.

### Cell cycle analysis

SUPM2 parental, SUPM2 LR1000 resistant to lorlatinib, Jeko-1 and Maver cells were treated with 1 µM concentration of VCB1 or BMS for 48 h. After the treatment, cells were fixed in 70% ice-cold ethanol, and PI staining was performed using the FxCycle PI/RNase Staining Solution (Thermo Fisher Scientific). Data from the gated tumor cells were acquired by running the samples on a FACSCanto II instrument. Cell cycle plots of quiescent (G0-phase) and actively cycling tumor cells (G1-, S-, G2M-phase) were generated with FlowJo software.

### Metabolomics Analysis

Metabolomics analyses were conducted at The Wistar Institute Proteomics and Metabolomics Shared Resource with slight modifications as previously described(35–37). The Global metabolomic analysis was performed on SUPM2 and SUPM2-LR1000 (SLR, lorlatinib resistant) cell lines were cultured in 5% dialyzed FBS and RPMI media and treated with 1.0 µM ACLYi VCB1 or BMS for 24 hr. The cells were spun down at 1000 rom and washed with cold PBS and snap frozen in liquid nitrogen. The pellets were sent to Wistar Metabolomics core.

### Global Lipidomics Analysis

The Global Lipidomics analysis was performed as described earlier with modification to our cell lines(38). SUPM2 and SUPM2-LR1000 (SLR, lorlatinib resistant) cell lines were cultured in 5% dialyzed FBS and RPMI media and treated with 1.0 µM ACLYi VCB1 or BMS for 24 hr. The cells were spun down at 1000 rpm and washed with cold PBS and snap frozen in liquid nitrogen. The pellets were sent to Wistar Metabolomics core. Samples were spiked with EquiSplash LIPIDOMIX internal standards (Avanti Polar Lipids), and lipids were extracted using a methyl tert-butyl ether (MTBE)-based method. The upper organic phase was collected, dried, and reconstituted in 9:1 methanol before storage at −20°C. Lipid extracts (10 µL per injection) were analyzed in randomized order by LC–MS/MS using a Thermo Scientific Q Exactive HF-X mass spectrometer coupled to a Vanquish UHPLC system. A pooled quality-control (QC) sample was analyzed periodically throughout the sequence to monitor analytical performance. Data were acquired separately in positive- and negative-ion modes following reversed-phase chromatographic separation on a C30 column. Lipid species were identified by MS/MS fragmentation using Thermo Scientific LipidSearch 4.2 and quantified based on peak areas. Identification confidence, expected adducts, retention-time windows, peak quality, QC reproducibility, and data completeness were used for quality filtering. Deuterated internal standards, species outside expected retention-time windows, lipids with average peak quality <0.7, QC coefficient of variation >30%, and species with fewer than 30/30 valid experimental measurements were excluded. Relative lipid abundances were corrected using EquiSplash standards for the corresponding lipid classes and quantified using TraceFinder 4.1. Where appropriate, data were additionally normalized to total lipid signal or sample weight.For lipid species-level comparisons, log2-transformed data were analyzed using Student’s *t*-test, with Benjamini–Hochberg correction for multiple testing. Differentially abundant lipids were defined as those exhibiting ≥1.5-fold change with *p*<0.05 or *q*<0.05. Lipid class abundances were calculated by summing peak areas of individual species within each class and were compared using Student’s *t*-test. MS/MS analysis did not resolve acyl-chain positional isomers or double-bond positions.

### Statistical analysis

Statistical analysis was performed via GraphPad Prism 8. Quantitative data are presented as mean ± standard deviation (SD) and all statistical tests were performed with a minimum of three biological replicates. If not otherwise stated, statistical testing was conducted by a two-sided unpaired t test to determine significant differences between the control and corresponding treatment conditions. A p-value of < 0.05 was considered statistically significant (*, p <0.05; **, p < 0.01; ***, p < 0.001; ****, p < 0.0001).

### Data availability

We confirm that all relevant data and methods are included in the main article, and the supplementary material of RNA-seq, Metabolomics and lipidomics will be uploaded to the appropriate website.

## Results

### Structure-based identification and validation of a novel ACLY inhibitor

ACLY is a central enzyme linking citrate metabolism to acetyl-CoA production and downstream lipid biosynthesis. We sought to identify novel small-molecule ACLY inhibitors using a structure-based drug discovery approach as shown in silico screening schematic (Fig.1A). The human ACLY crystal structure was utilized to perform both ligand-based and structure-based in silico screening of the National Cancer Institute (NCI) chemical database to identify potential inhibitors targeting the ATP-binding site within the ATP-grasp domain of ACLY as shown in the schematic (Fig.1B). To identify novel ATP-citrate lyase (ACLY) inhibitors, a structure-based pharmacophore model was generated from the adenine nucleoside-binding interactions of ADP within the human ACLY binding site (PDBID: 3PFF) as shown (Fig.1C). The resulting five-point 3D pharmacophore comprised one hydrogen-bond donor (Don), three hydrogen-bond acceptors (Acc), and one aromatic ring center (Ar), reflecting the key interactions responsible for ligand recognition (Fig.1D). To identify the docking pocket of ATP binding pocket, we used the “Site finder” module of MOE (Chemical Computing Group, Montreal, Canada) to identify the potential pocket and filled with dummy atoms. The dummy atom locations in the ATP binding site of ACLY structure were used to dock the matched and energetically favorable conformers. Location of Dummy “atoms” (sphere) identified in the ATP binding pocket. Red sphere has contact with atom(s) of ACLY protein and gray sphere does not have contact with ACLY protein. The amino acid residues forming the pocket are shown by gray for carbon, red for oxygen and blue for nitrogen (Fig.1D). The docking of active ACLY inhibitor VCB#1(magenta surface) into the ATP binding pocket of ACLY (khaki colored ribbon) and the amino acids in ACLY lining the binding pocket of VCB11 are shown by cornflower blue for carbon; red for oxygen and blue for nitrogen (Fig.1E). The docked VCB1 in the ATP binding site of ACLY forms five favorable hydrogen bonds with the side chain atoms of amino acids of ACLY protein R66, E108, E201 and D216. Two R66 forms two hydrogen bonds with the two nitrogens in the six-member ring as shown (Fig.1F) Distance constraints between pharmacophoric features were incorporated to preserve the spatial arrangement observed in the human ACLY–ADP crystal structure. The model accurately mapped the critical binding determinants of the adenine-binding pocket and was subsequently employed as a query for virtual screening to identify compounds with the potential to engage the ATP-binding site of ACLY. Our 3D pharmacophore search as described lead to 131 unique compounds. Based on the MOE docking scoring of each conformation-ACLY complex, the rank ordering of the complexes was carried out, we selected the top 100 ACLY compound conformation complexes for further reduced to a list of unique compounds. Based on the availability in NCI repository, we received 11 compounds for ACLY activity screening assay. The 2D chemical structure of three active compounds and reference compound are shown in (Fig.1G). The other selected compound 2D chemical structures are shown (Supplementary Fig.1). Next, the three active compounds were subsequently tested for their ability to inhibit recombinant human ACLY enzymatic activity, using the previously reported ACLY inhibitor BMS-303141 as a reference compound. The results showed that our lead compounds compound#1 (VCB1), compound#2 (VCB2) and compound#7 (VCB7) demonstrated an IC₅₀ of 0.91, 0.46 and 1.19 µM respectively, confirming direct inhibitory activity against recombinant ACLY (Fig.1H). Whereas the reference compound BMS-303141 inhibited recombinant human ACLY with an IC₅₀ of 0.26 µM in our study.

**Figure 1.**
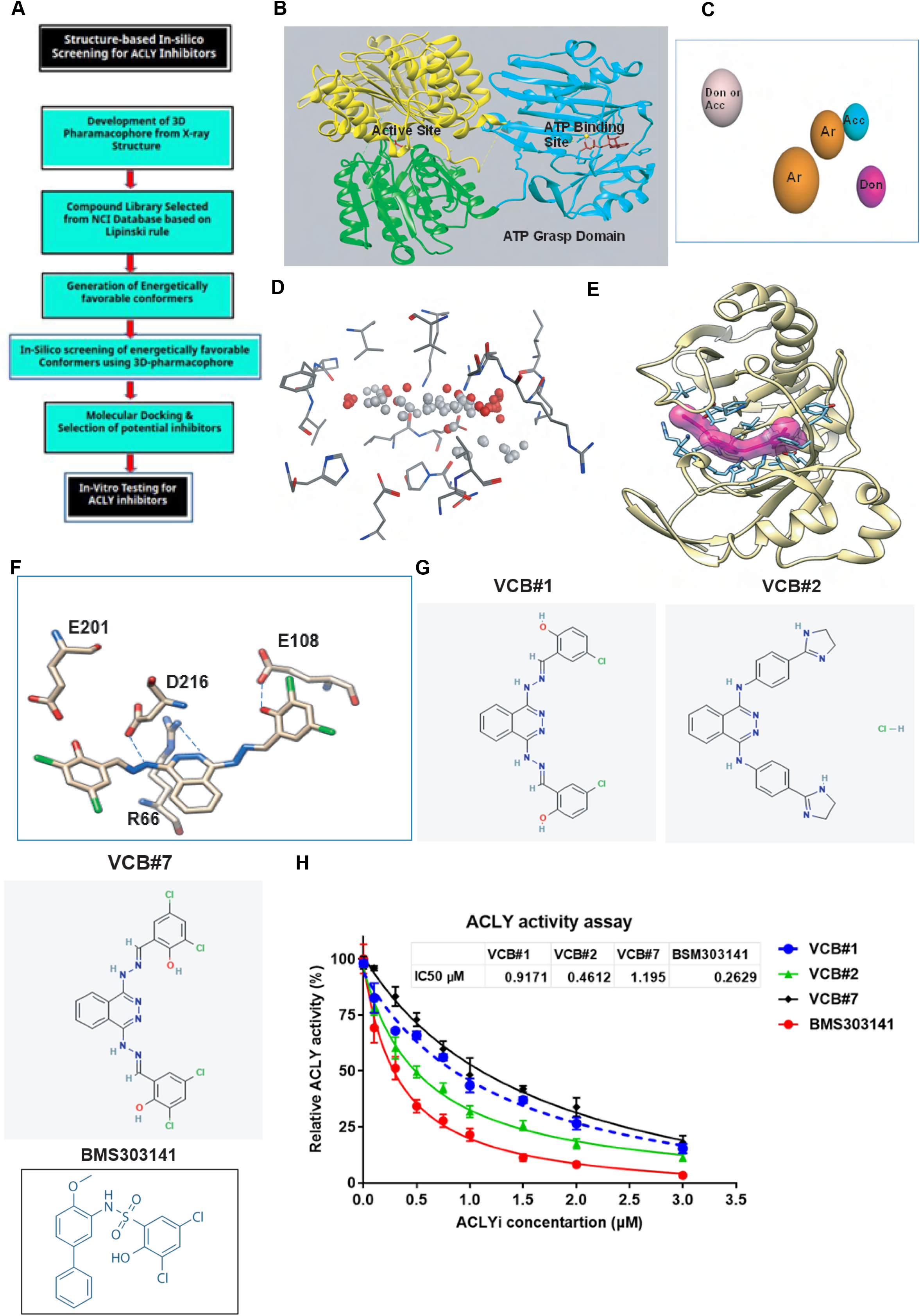
Structure-based identification and biochemical validation of novel ACLY inhibitors. (A) Schematic overview of the structure-based drug discovery workflow used to identify novel small-molecule inhibitors targeting the ATP-binding site of human ACLY. (B) Generation of a structure-based pharmacophore model using the adenine nucleoside-binding interactions of ADP in the human ACLY crystal structure (PDB: 3PFF). (C) Five-point 3D pharmacophore model comprising one hydrogen-bond donor (Don), three hydrogen-bond acceptors (Acc), and one aromatic ring center (Ar), with spatial distance constraints derived from the ACLY–ADP structure. (D) Identification of the putative ATP-binding pocket using the MOE Site Finder module; dummy atoms within the pocket are shown as spheres, with protein-contacting and non-contacting sites indicated separately. (E) Docking pose of VCB1 within the ATP-binding pocket of human ACLY, showing the compound and surrounding pocket residues. (F) Predicted molecular interactions between VCB1 and ACLY, showing five favorable hydrogen bonds involving residues R66, E108, E201, and D216, including two interactions formed by R66 with nitrogen atoms of VCB1. (G) Chemical structures of the three active compounds identified from the NCI chemical database together with the reference ACLY inhibitor BMS-303141. (H) Biochemical validation of ACLY inhibition using recombinant human ACLY. BMS-303141 inhibited ACLY with an IC₅₀ of approximately 250 nM, whereas VCB1 inhibited ACLY with an IC₅₀ of approximately 560 nM, confirming direct enzymatic inhibition. Overall, the workflow identified VCB1 as a novel ATP-site-directed ACLY inhibitor for Furthermore cellular and mechanistic characterization.

### Genetic silencing of ACLY in ALK+ALCL cell line and xenograft model revealed ACLY as a therapeutic target

To investigate the requirement for ACLY in ALK-positive ALCL tumor growth, we initially focused on SU-DHL-1 cells, which showed relatively high ACLY expression among ALK-positive ALCL cell lines as confirmed in our previous study(6). ACLY is a central metabolic regulator implicated in tumor growth across multiple malignancies, and genetic ACLY silencing has been shown to induce growth arrest and apoptosis in cancer cells(2, 39). However, genetic validation of *ACLY* dependence in lymphoma xenograft models remains comparatively limited. We therefore generated SUDHL-1 cells expressing control shRNA (shControl) or *ACLY*-targeting shRNA (shACLY) and evaluated tumor formation following subcutaneous flank injection. All six mice injected with shControl cells developed tumors and detectable tumors observed 64 days after implantation and subsequently exhibited rapid tumor growth (Supplementary Fig.2A). Mean tumor volume increased from 291.8 ± 97.7 mm³ on day 64 to 1271.9 ± 521.5 mm³ by day 76 (n=6) (Supplementary Fig.2B). At the experimental endpoint, excised shControl tumors weighed 0.6–3.2 g, with a mean weight of 1.82 ± 0.99 g (Supplementary Fig.2C). In contrast, no palpable tumors were detected in seven mice injected with shACLY cells throughout the observation period, indicating that ACLY depletion markedly impaired detectable tumor establishment in vivo (Supplementary Fig.2D). Together with recent genetic evidence of ACLY dependence in T-cell acute lymphoblastic leukemia(40). These findings provided further support for investigating ACLY as a potential therapeutic target in ALK-positive ALCL.

**Figure 2.**
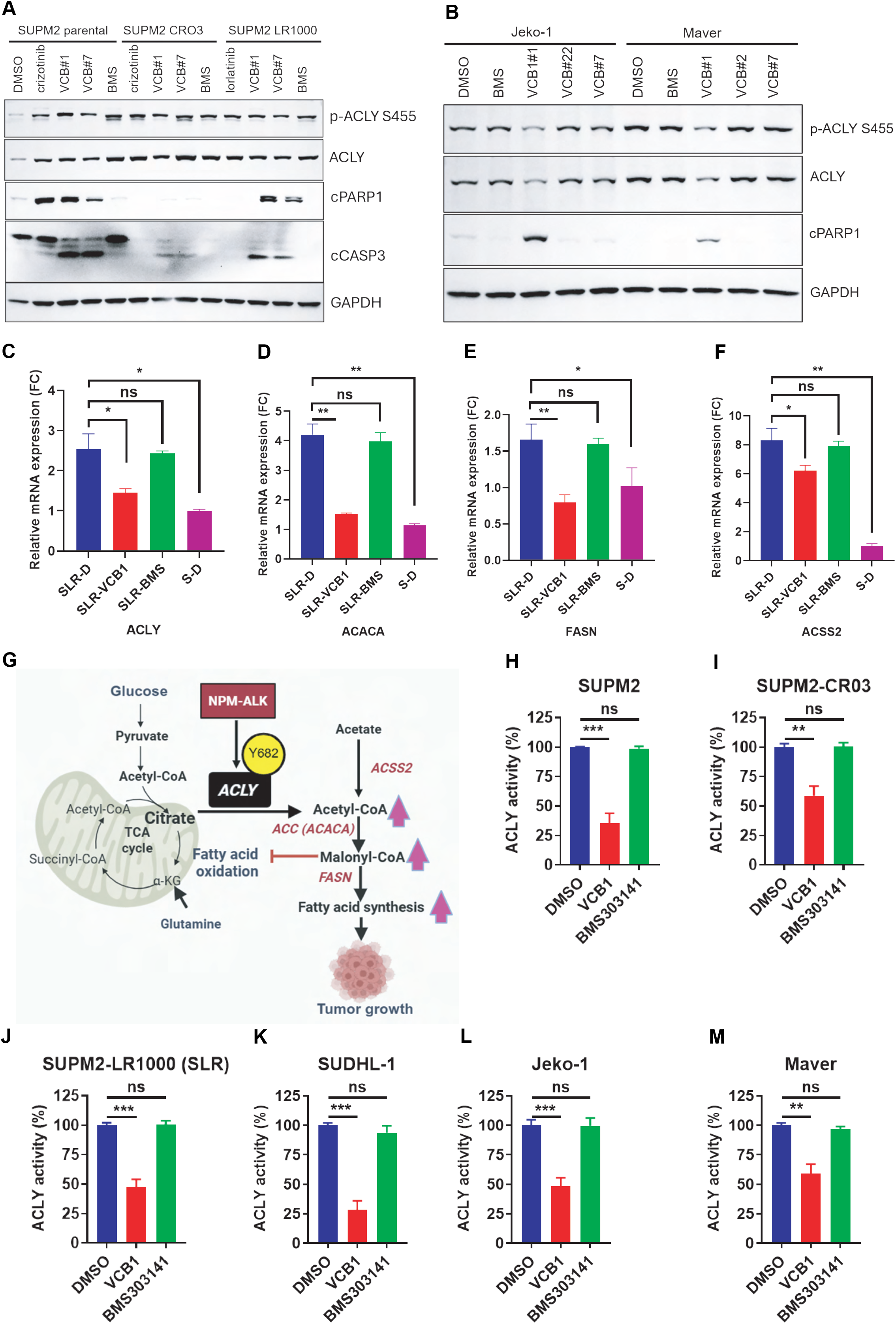
Novel ACLY inhibitor decreases ACLY activity and promotes cell death. (A) SUPM2 parental-ALKi sensitive and SUPM2-LR1000 ALKi-resistant ALK-positive ALCL cells were treated with ACLY inhibitors VCB1, VCB2, VCB7, or BMS-303141 (BMS) at 1.0 μM for 48 h in 5% FBS medium. Cell lysates were analyzed by immunoblotting for cleaved PARP1 and cleaved caspase-3 as markers of apoptosis. (B) Ibrutinib-sensitive Jeko-1 and ibrutinib-resistant Maver MCL cells were treated with the indicated ACLY inhibitors under the same conditions, and apoptosis was assessed by immunoblotting for cleaved PARP1 and cleaved caspase-3. (C–F) TKI-resistant lymphoma cells were treated with VCB1 or BMS at 1.0 μM for 24 h, followed by analysis of ACLY, ACACA, FASN, and ACSS2 expression. (G) Schematic representation of the proposed NPM-ALK–ACLY metabolic axis, in which NPM-ALK-mediated phosphorylation of ACLY at Y682 promotes ACLY activity and supports de novo fatty-acid synthesis. (H–M) Cellular ACLY activity was measured following treatment with VCB1 or BMS in S-sensitive and SLR-resistant ALK-positive ALCL cells and in MCL cell lines. Data are representative of independent experiments. Statistical significance was determined using the appropriate statistical test; P < 0.05 was considered significant.

### A novel ACLY inhibitor, VCB1 suppresses ACLY activity and promotes cell death

Our xenograft study provided a rationale for pharmacologic targeting of ACLY in ALK-positive ALCL. We therefore next evaluated three lead ACLY inhibitors, VCB1, VCB2, and VCB7, together with the reference ACLY inhibitor BMS-303141 (BMS), using the paired SUP-M2 parental and lorlatinib-resistant SUP-M2-LR1000 cell lines. To evaluate the cellular activity of our lead ACLY inhibitors (VCB1, VCB2, and VCB7) along with the reference inhibitor BMS, we examined their effects on lymphoma cell viability and apoptotic signaling. ALK inhibitor (ALKi)-sensitive SUPM2 parental (S) cells, as well as crizotinib- and lorlatinib-resistant lines (SUPM2 CR03 and SUPM2-LR1000 [SLR]), were treated with 1.0 μM of each compound in medium containing 5% FBS for 48 hours. Immunoblot analysis revealed that among the three novel compounds, VCB1 and VCB7 induced a marked increase in cleaved PARP1 and cleaved caspase-3, indicating the activation of apoptotic cell death (Fig. 2A). Notably, this apoptotic response was most pronounced in the parental SUPM2 cells, followed by the lorlatinib-resistant SLR cells, while the crizotinib-resistant line exhibited the least sensitivity. We next investigated whether the activity of these lead compounds extended to mantle cell lymphoma (MCL) models with differential sensitivity to the BTK inhibitor ibrutinib. Ibrutinib-sensitive Jeko-1 and ibrutinib-resistant Maver cells were treated under the same experimental conditions. Consistent with our observations in ALK+ ALCL, VCB1 induced robust cleavage of PARP1 in both MCL cell lines, whereas VCB2, VCB7 and the reference compound BMS-303141 elicited minimal apoptotic activation (Fig. 2B). These results demonstrate that the cytotoxic activity of the ACLY inhibitor VCB1 extends across genetically and therapeutically distinct lymphoma models. Based on these collective findings, VCB1 was selected as the primary lead compound for subsequent mechanistic studies and direct comparison with BMS-303141. We next investigated whether VCB1 altered ACLY expression and the expression of key metabolic enzymes associated with the ACLY-dependent lipogenic pathway. SLR cells were treated with VCB1 or BMS at 1.0 μM for 24 h, followed by RNA isolation and gene-expression analysis. VCB1 significantly reduced the expression of *ACLY* as well as the downstream lipogenic enzymes *ACACA* and *FASN* and the acetate-utilizing enzyme *ACSS2* (Fig. 2C–F). In contrast, BMS did not produce a comparable reduction in these metabolic genes. These findings suggested that VCB1 produces a broader suppression of the *ACLY*-dependent metabolic transcriptional program than the reference compound. Our previous studies demonstrated that NPM-ALK directly phosphorylates ACLY at Y682, thereby enhancing ACLY activity and supporting lipid biosynthesis in ALK+ lymphoma cells. Thus, ACLY represents a metabolic effector downstream of oncogenic NPM-ALK signaling, linking tyrosine kinase signaling to de novo lipid synthesis (Fig. 2G). To determine whether the cellular effects of VCB1 were associated with inhibition of ACLY enzymatic activity, we measured cellular ACLY activity following treatment with VCB1 or BMS in sensitive (SUPM2 parental and SUDHL1) and ALKi-resistant ALK+ ALCL and in MCL cell lines. VCB1 treatment produced a substantial reduction in cellular ACLY activity compared with BMS across the lymphoma models examined (Fig. 2H–M). Collectively, these results demonstrate that VCB1 inhibits ACLY activity in lymphoma cells, suppresses expression of key lipogenic enzymes, and induces apoptotic cell death. The substantially greater cellular activity of VCB1 compared with VCB2, VCB7 and BMS provided a rationale for selecting VCB1 as the lead compound for subsequent transcriptomic, metabolomic, and lipidomic studies.

### ACLY inhibition suppresses lipogenesis and induces metabolic and proliferative reprogramming

To define the transcriptional consequences of ACLY inhibition, we performed RNA sequencing (RNA-seq) in SUPM2 parental and SLR cells following treatment with VCB1 or BMS. Cells were treated with 1.0 μM compound for 24 h, and RNA from three biological replicates per condition was subjected to RNA-seq. The distribution of differentially expressed genes (DEGs) across the comparisons is summarized (Supplementary Fig. 3A). VCB1 induced a substantially stronger transcriptional response than BMS in both cell lines. The largest transcriptional response occurred in S-VCB1 versus S-DMSO cells, whereas S-BMS versus S-DMSO produced relatively few DEGs (Supplementary Fig. 3B–E). A similar pattern was observed in SLR cells, in which VCB1 produced substantially more transcriptional changes than BMS. Baseline comparison of untreated SLR and parental cells also demonstrated extensive transcriptional reprogramming associated with acquired ALKi resistance (Supplementary Fig. 3F). Using a threshold of log2 FC ≥1 and P<0.05, VCB1 treatment in parental S cells resulted in 793 upregulated and 209 downregulated genes, whereas 24,781 genes remained unchanged (Supplementary Fig. 3D). In comparison, BMS treatment resulted in only 42 upregulated and 65 downregulated genes, with 25,435 genes unchanged (Supplementary Fig. 2E). Many genes uniquely altered by BMS were non-coding RNAs, pseudogenes, or poorly characterized loci, whereas VCB1 produced a broader response involving protein-coding genes.

The more pronounced transcriptional response to VCB1 was retained in SLR cells, demonstrating that VCB1 engages a strong cellular response despite acquired ALKi resistance. Baseline comparison of untreated SLR and S cells identified 3,318 upregulated and 2,979 downregulated genes in SLR cells, with 21,183 genes remaining unchanged. Thus, acquired ALKi resistance was associated with extensive transcriptional reprogramming, creating a metabolically distinct cellular state in which ACLY inhibition produced a broad adaptive and stress response. To capture moderate but potentially biologically important transcriptional changes, we subsequently applied a threshold of log2 FC ≥0.5 and P<0.05. This analysis identified 1,427 DEGs in TKI-sensitive cells and 3,712 DEGs in TKI-resistant SLR cells. Venn analysis identified 561 genes unique to S cells, 2,839 unique to SLR cells, and 866 commonly regulated genes (Fig. 3A). Thus, the transcriptional consequences of VCB1 were substantially broader in resistant cells, consistent with greater metabolic and stress adaptation in the resistant state. To identify molecular pathways affected by VCB1 mediated ACLY inhibition, a pathway enrichment analysis was carried out using the Database for Annotation, Visualization and Integrated Discovery (DAVID) focusing on the Gene Ontology Biological Processes (GO_BP), Kyoto Encyclopedia of Genes and Genomes (KEGG), Reactome, and Wiki Pathways(41). KEGG pathway analysis using NIH-DAVID demonstrated enrichment of ribosome biogenesis, amino-acid biosynthesis, HIF-1 signaling, TNF signaling, carbon metabolism, and broader metabolic pathways (Fig. 3B, C). Among the 866 genes commonly regulated by VCB1 in sensitive and resistant cells, VCB1 suppressed key components of the lipogenic program, including *ACLY, ACACA, FASN, GPAM, and HACD2*, together with genes involved in lipid remodeling and homeostasis, including *INSIG1, INSIG2, PLIN2, STARD4,* and *SLC27A4*. These changes were consistent with suppression of ACLY-dependent de novo lipid synthesis. VCB1 simultaneously altered genes involved in glycolysis, hypoxia, and metabolic stress, including *SLC2A1, SLC2A3, PFKFB3, PFKFB4, PGK1, CA9, BNIP3, BNIP3L*, and *DDIT4. PFKFB4* was induced in both cellular states, with a stronger induction in sensitive cells, consistent with a compensatory response to altered glycolytic and energetic demands. In parallel, genes involved in cell-cycle progression and mitosis, including *CCNB1, CDC20, PLK1, CENPE, CENPF, KIF20A, KIF23*, and *PTTG1*, were predominantly downregulated (Fig. 3D–F). These findings indicate that ACLY inhibition couples metabolic stress to suppression of proliferative programs. The response was more pronounced in SLR cells, whereas VCB1 suppressed genes involved in fatty-acid synthesis, elongation, cholesterol metabolism, and lipid remodeling, including *FASN, ACACA, HACD2, HACD3, LPCAT3, LPCAT4, DHCR24, MECR, MCAT, CPT1A*, and *PPARGC1B. PPARGC1B*, which regulates oxidative and lipid metabolic programs, was particularly reduced in resistant cells. In contrast, genes associated with fatty-acid oxidation, lipid uptake, and lipoprotein metabolism, including *ACADS, ACADVL, PPARA, LPL, LRP1, LDLR, VLDLR, HMGCL*, and *OSBPL5*, were increased (Fig. 3G). Thus, ACLY inhibition suppressed endogenous lipid synthesis while inducing a compensatory transcriptional program favoring lipid uptake and oxidative metabolism. Basal comparison of S and SLR cells revealed further metabolic differences. SLR cells preferentially expressed genes involved in cholesterol biosynthesis, lipid storage, remodeling, and uptake, including *DHCR24, FDFT1, MSMO1, FADS2, DGAT2, MOGAT2, APOC1, APOE, LIPE, HILPDA, and LDLRAP1*, whereas parental S cells showed greater expression of genes involved in glycerophospholipid synthesis and remodeling (Fig. 3H,I). Examination of SREBP-associated genes further supported suppression of lipid biosynthesis. VCB1 reduced *HMGCR, GPAM, SLC27A4, ACLY, DHCR7, ACACA, HMGCS1, FASN,* and *PPARGC1A*, while increasing *OSBPL3, OSBPL6, INSIG1, INSIG2, DHCR24, HMGCL*, and *LRP1* (Fig. 3J). Next, we performed STRING-protein analysis and results demonstrated an interconnected ACLY-associated metabolic network (Fig. 3K). VCB1 also increased expression of cell-cycle checkpoint and stress-response genes, including *CDKN1A, E2F7, SESN2, DDIT4, BNIP3,* and *BTG1*, while reducing mitotic regulators such as *CDC20 and KIF20A* (Fig. 3L). To further, STRING-protein pathway analysis revealed enrichment of DEGs involved in TNFα and IL-17 related genes (Fig.3M).

**Figure 3.**
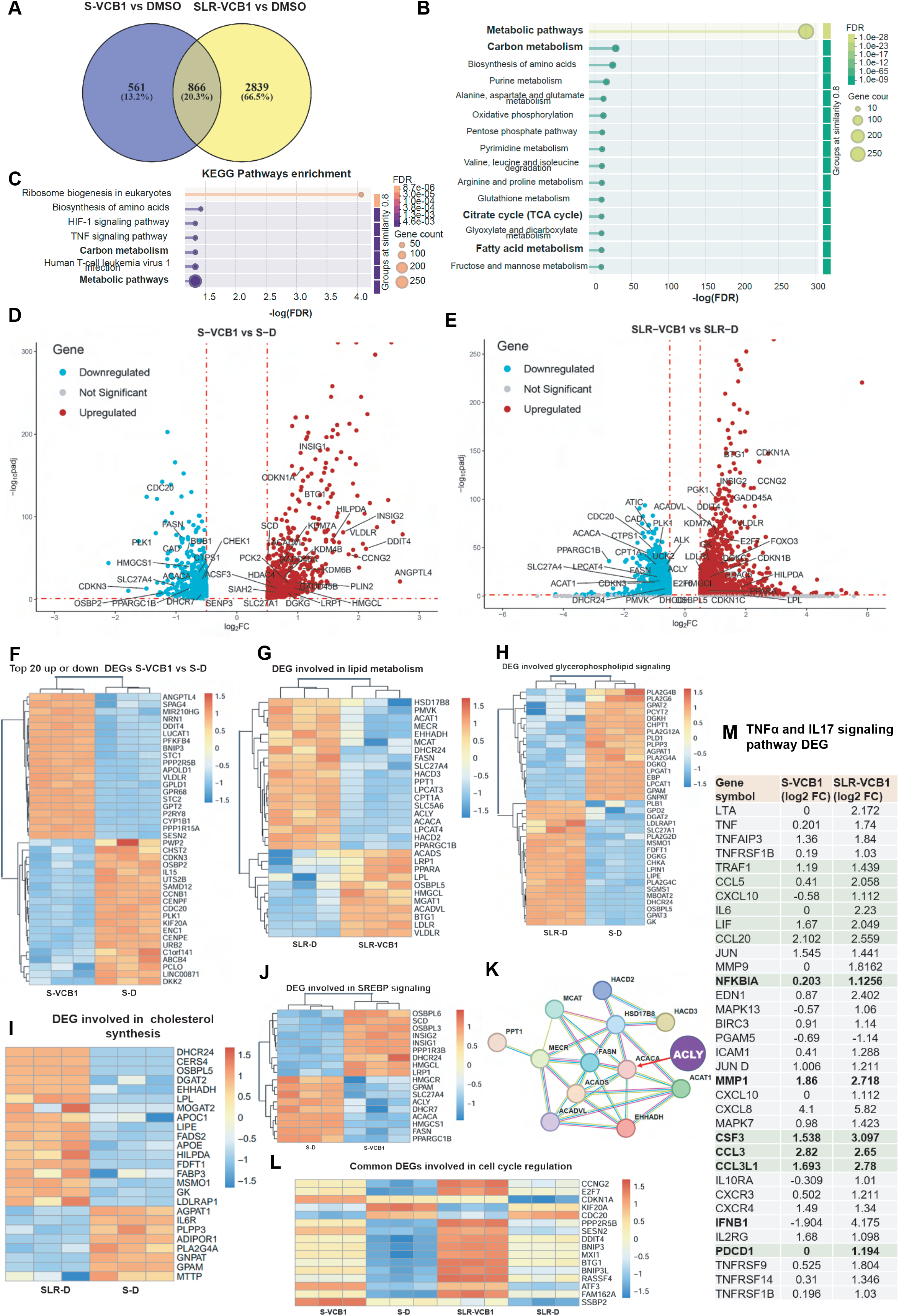
RNA-sequencing reveals metabolic, proliferative, and inflammatory reprogramming following ACLY inhibition. (A) Number of differentially expressed genes (DEGs) identified in TKI-sensitive (S-VCB1) and TKI-resistant (SLR-VCB1) cells following VCB1 treatment compared with DMSO controls. DEGs were defined using log2 FC ≥ 0.5 and P < 0.05. Venn diagram shows DEGs unique to each cellular context and genes commonly regulated by VCB1. (B) KEGG pathway enrichment analysis of DEGs identified following ACLY inhibition using NIH-DAVID, showing enrichment of ribosome biogenesis, amino-acid biosynthesis, HIF-1 and TNF signaling, carbon metabolism, and other metabolic pathways. (C) Distribution of metabolism-associated DEGs across major metabolic categories, highlighting carbon metabolism as a prominent transcriptionally affected pathway. (D,E) Volcano plots showing the transcriptional response to VCB1 in S-sensitive and SLR-resistant cells (F) Heatmap of representative DEGs in S-sensitive cells following VCB1 treatment. (G) Heatmap of lipid-metabolism-associated DEGs in SLR-resistant cells following VCB1 treatment. (H) Basal glycerophospholipid metabolism-associated gene expression in S-sensitive versus SLR-resistant cells, demonstrating distinct lipid synthesis and remodeling programs associated with S sensitivity and resistance. (I) Basal cholesterol- and lipid-metabolism-associated gene expression in parental SUP-M2 and SLR-resistant SLR-D cells. (J) SREBP-associated DEGs following VCB1 treatment in S-sensitive cells. (K) In schematic display showing ACLY networking genes by suing strin-protein interactome website. (L) Cell-cycle-associated DEGs following ACLY inhibition, showing induction of checkpoint and stress-response genes and suppression of mitotic regulators. (M) DEGs associated with TNF and IL-17 signaling pathways following ACLY inhibition in S-sensitive and SLR-resistant cells.

To better visualize the interactions among the common differentially expressed genes (DEGs) across both cell lines, we performed a network analysis using the STRING database. First, we mapped all common DEGs meeting the threshold of log2 fold change (FC) +/- 1.0 and (P < 0.05), irrespective of their functional classification (Supplementary Fig. 4). Using the same statistical criteria, we specifically isolated and mapped the network of common DEGs involved in metabolic pathways (Supplementary Fig. 5A). Because ACLY inhibition alters lipid levels and remodels gene expression in response to lipid-mediated stress, we next performed a Gene Ontology (GO) biological process analysis. This analysis revealed a significant enrichment of DEGs involved in the cellular response to lipids (Supplementary Fig. 5B). Furthermore, network analysis highlighted distinct signaling clusters related to both TNF and HIF-1 signaling pathways (Supplementary Fig. 5C and D).

### VCB1 induces DEGs involved in Metabolic and transcriptional reprogramming in TKI-sensitive and TKI-resistant cells

To define metabolic function associated with DEGs, cells were analyzed using a log2 fold-change threshold of ≥ +0.5 or ≤ −0.5. RNA-seq identified a distinct set of common metabolic DEGs shared between two cell lines (Table-1). These genes revealed coordinated alterations in glycolysis, lipid metabolism, mitochondrial function, and amino-acid metabolism, including induction of glycolytic genes such as PFKFB4 and PFKL, together with suppression of key lipogenic and mitochondrial genes including *ACACA, FASN, CS, DLAT*, and *PPARGC1B*. Notably, *PPARGC1B* suppression was substantially greater in resistant cells, suggesting enhanced impairment of mitochondrial oxidative and lipid metabolic programs during TKI resistance. In contrast, SUPM2 parental, specific DEGs displayed a metabolic signature characterized by increased glycolysis, lipid uptake/storage, amino-acid biosynthesis, and nutrient-stress signaling, exemplified by *PFKFB4, ANGPTL4, VLDLR, HILPDA, GPT2, ASNS, SESN2*, and *DDIT4* (Supplementary Table-1). These changes suggest that TKI-sensitive cells respond to metabolic perturbation by activating alternative nutrient-utilization and metabolic stress-adaptation pathways. The SLR specific DEGs revealed a broader and more complex metabolic phenotype, characterized by enhanced glucose uptake and glycolytic adaptation, glutamine metabolism, redox/glutathione metabolism, and stress responses, including induction of *SLC2A1, PDK1, PDK3, GLUL, CHAC1*, and *SOD2*. At the same time, resistant cells showed extensive suppression of mitochondrial oxidative metabolism, TCA-cycle activity, fatty-acid metabolism, phospholipid synthesis, and nucleotide biosynthesis, including *CAD, CPT1A, IDH3A, OGDH, FH, DHODH*, and multiple mitochondrial respiratory-chain genes (Supplementary Table-2). Collectively, the shared and group-specific metabolic DEGs define distinct metabolic signatures associated with TKI sensitivity versus resistance. While both populations exhibit suppression of key lipogenic and mitochondrial pathways, TKI-sensitive cells preferentially activate metabolic stress, nutrient uptake, and biosynthetic adaptation, whereas TKI-resistant cells display a more extensive metabolic reprogramming characterized by glycolytic dependence, glutamine/redox adaptation, and broad mitochondrial and anabolic metabolic suppression.

### VCB1 treatment and cancer hallmark analysis on DEGs revealed metabolic reprogramming and inflammatory adaptation

Analysis of cell-line-specific DEGs revealed distinct biological responses to ACLY inhibition. At the log2 FC ≥0.3 threshold, VCB1-treated SLR cells showed enrichment of cancer hallmarks associated with genome instability, reprogramming energy metabolism, replicative immortality, and resistance to cell death (Fig. 4A). When downregulated genes were analyzed separately at the log2 FC ≥0.7 threshold, reprogramming energy metabolism was significantly enriched, involving 27 genes (P=0.00257; FDR-adjusted P=0.0231) (Fig. 4B). The enriched genes included *ACLY, ACACA, FASN, PPARG, PFKM, IDH1, IDH3A, CS,* and *DLAT*, together with mitochondrial respiratory-chain and TCA-cycle genes. These findings indicate that VCB1 suppresses a broad tumor-associated energy-metabolic program rather than affecting lipid synthesis alone. In contrast, upregulated DEGs in SLR cells were enriched for the sustained angiogenesis hallmark (Fig. 4C). At a log2 FC threshold of 0.7, the strongest enrichment was driven by *TNF, THBS1, TGFBR1, CXCR3, MMP1, ITGA5, IL6, SDC4, CSF3*, and *CXCL10*. Tumor-promoting inflammation and immune-evasion programs were also represented by genes including *TNF, TNFSF13B, PDCD1, IL6, LY96, FCGR1A, CD247, IL2RG, CXCL10*, and *BCL6*. These data indicate that ACLY inhibition elicits an inflammatory and immune-associated transcriptional response in addition to metabolic suppression. Analysis of genes commonly upregulated in S-VCB1 and SLR-VCB1 cells at log2 FC ≥0.7 revealed enrichment for sustained angiogenesis, resistance to cell death, and reprogrammed energy metabolism (Fig. 4D). Similar analysis using a log2 FC ≥0.5 cutoff demonstrated enrichment of reprogramming energy metabolism and sustained angiogenesis among common DEGs (Supplementary Fig. 6A, B). Common upregulated genes were additionally enriched for sustained angiogenesis, sustained proliferative signaling, resistance to cell death, and reprogrammed energy metabolism, whereas downregulated genes were significantly enriched for reprogrammed energy metabolism (Supplementary Fig. 6C–F). Collectively, these analyses demonstrate that VCB1 produces a conserved metabolic response across TKI-sensitive and TKI-resistant cells, while the resistant state additionally engages stronger inflammatory, angiogenic, and adaptive stress programs.

**Figure 4.**
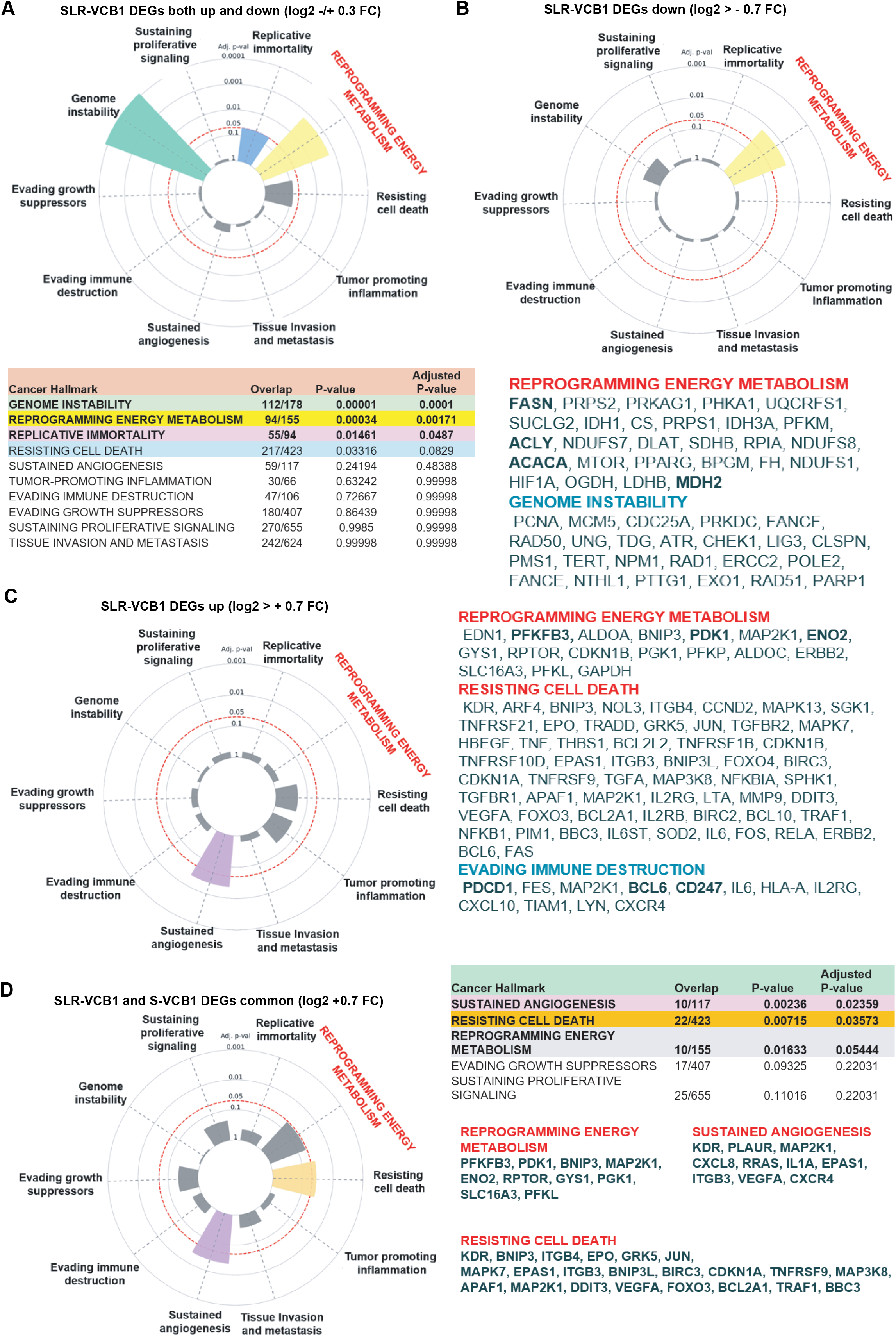
Cancer hallmark enrichment of common and cell-line-specific DEGs following ACLY inhibition. (A) Cancer hallmark enrichment analysis of DEGs unique to SUPM2-LR1000 (SLR) cells following VCB1 treatment using a log2 fold-change cutoff of ≥0.3 up and down regulated. The cancer hallmark pathways are shown on the table below. (B) Hallmark enrichment analysis of downregulated DEGs in SLR-VCB1 cells using log2 fold-change of -0.7 down regulated. The significant DEGs related to each hallmark are shown below. (C) Hallmark analysis of upregulated DEGs in SLR-VCB1 cells demonstrating significant enrichment of sustained angiogenesis. The corresponding enriched genes are shown in the right panel. (D) Hallmark enrichment analysis of commonly upregulated DEGs shared between S-VCB1 and SLR-VCB1 cells using a log2 fold-change cutoff of ≥0.7. Sustained angiogenesis, resisting cell death, and reprogramming energy metabolism were significantly enriched, with the corresponding genes shown in the right panel.

### VCB1 induces extensive metabolic remodeling characterized by citrate accumulation and nucleotide depletion

To determine whether the transcriptional consequences of ACLY inhibition were accompanied by metabolic changes, we performed integrated untargeted metabolomic analysis. Untargeted metabolomics of TKI-sensitive SUPM2 cells treated with DMSO, VCB1, or BMS identified 307 metabolites. Significance was defined as a ≥1.5-fold change with adjusted q<0.05. Hierarchical clustering demonstrated a distinct metabolic profile in VCB1-treated cells compared with DMSO controls (Fig. 5A). A stringent fourfold cutoff was subsequently used to highlight the most strongly altered metabolites (Fig. 5B). VCB1 markedly depleted metabolites associated with purine and pyrimidine metabolism. The most strongly decreased metabolites with significant q-value of < 0.05 included with fold change are dADP (−34.06-fold), cytosine (−29.36-fold), AMP (−20.61-fold), UMP (−16.26-fold), and TMP (−16.23-fold). These coordinated changes revealed a broad disruption of nucleotide metabolism following ACLY inhibition. VCB1 also induced marked accumulation of central-carbon metabolites, including D-myo-inositol 1,4-bisphosphate (1810.66-fold), DHAP/GAP (151.54- and 33.37-fold), phosphoenolpyruvate (99.59-fold), and 3- phosphoglyceric acid (20.93-fold), together with glucosamine-6-phosphate, mannose-6-phosphate, acetyl-CoA, and Asp-Asp. These findings indicate extensive remodeling of glycolysis and central carbon metabolism (Fig. 5B). Conversely, citrate, the direct substrate of ACLY, accumulated 2.64-fold following VCB1 treatment (q=0.0017). Although citrate did not reach the fourfold cutoff used for the focused heatmap, analysis of raw metabolite peak intensity confirmed its accumulation following VCB1 treatment but not in BMS treated cells (Fig. 5C). Citrate accumulation therefore provided an important metabolic signature consistent with intracellular inhibition of ACLY as clarified by the schematic conversion of citrate and acetyl-CoA via ACLY shown in Fig.5C lower panel. On the other hand, it is possible to notice the citrate in the volcano plots showing the deregulated metabolites (Fig.5D). Not surprisingly, we recorded an increase of acetyl-CoA following VCB1 treatment, because untargeted metabolomics does not resolve cytosolic, nuclear, and mitochondrial acetyl-CoA pools, and this finding does not necessarily indicate preservation of the cytosolic acetyl-CoA pool required for ACLY-dependent lipogenesis. Acetyl-CoA may also arise from acetate utilization through ACSS2(42, 43) or from mitochondrial fatty-acid oxidation(44). Consistent with this interpretation, ACLY loss has previously been shown to increase acetate dependence through ACSS2 in CD8 T cells(45). Thus, the increase in total acetyl-CoA may reflect compensatory production from alternative metabolic sources rather than maintenance of ACLY-derived cytosolic acetyl-CoA. On the other hand, BMS-303141 produced substantially fewer metabolic changes and did not induce comparable citrate accumulation (Fig. 5E). Next, we performed Pathway enrichment and pathway impact analysis by using MetaboAnalyst online tool as described earlier(35, 46). The results showed that significantly altered metabolites identified purine and pyrimidine metabolism as the most strongly enriched pathways (Fig. 5F). Pathway impact analysis further identified pyrimidine, purine, pyruvate, and citrate metabolism as major affected pathways (Fig. 5G). SLR cells contained increased abundance of multiple nucleotide metabolites, indicating a metabolically distinct state characterized by enhanced nucleotide availability (Fig. 5H). Furthermore, the ACLYi VCB1 cell line produced an overall metabolic response in resistant cells like that of the SUMP2 parental line (Fig. 5I). Together, these data demonstrate that VCB1mediated ACLY inhibition produces coordinated remodeling of central carbon and nucleotide metabolism.

**Figure 5.**
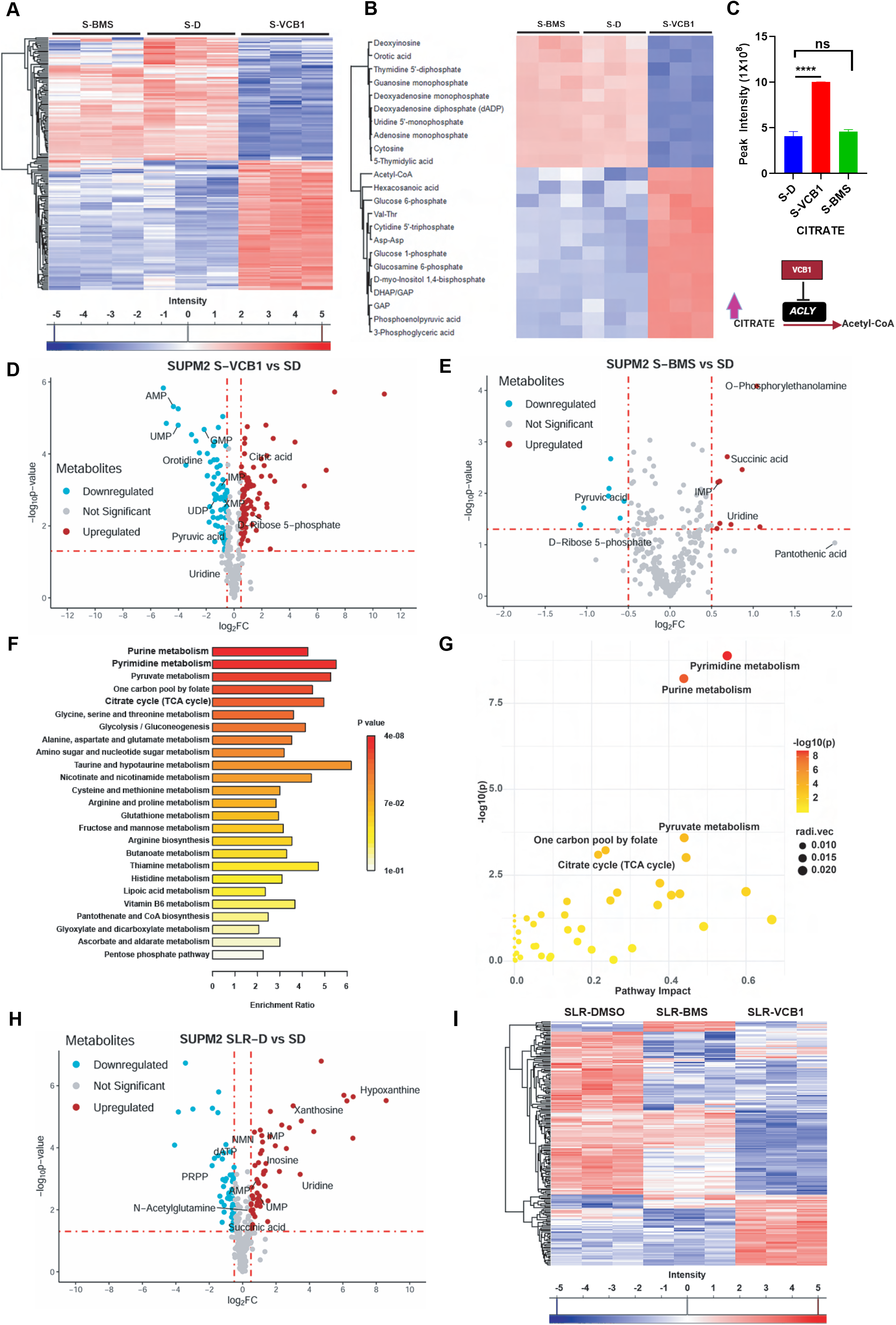
Untargeted metabolomics reveals distinct metabolic consequences of ACLY inhibition by VCB1. (A) Hierarchical clustering heatmap of significantly altered metabolites identified by untargeted metabolomics in TKI-sensitive cells (SUPM2) treated with DMSO, VCB1, or BMS-303141. Metabolites were considered significant at q<0.05 and a minimum 1.5-fold change. (B) Heatmap showing the most strongly altered metabolites using a 4-fold change cutoff, highlighting marked depletion of purine and pyrimidine-related metabolites and accumulation of glycolytic and central-carbon intermediates following VCB1 treatment. (C) Citrate abundance based on metabolomics peak abundance quantification in TKI-sensitive, SUPM2 cells treated with DMSO, VCB1, or BMS-303141, the schematic of Citrate to acetyl-CoA conversion via ACLY shown in bottom panel. (D) Volcano plot shows significantly altered metabolites following VCB1 treatment. (E) Volcano plot showing metabolic changes following BMS-303141 treatment, demonstrating substantially fewer alterations and no significant accumulation of citrate. (F) Metabolic pathway enrichment analysis of significantly altered metabolites following VCB1 treatment using MetaboAnalyst. (G) Pathway impact analysis showing the major metabolic pathways affected by VCB1, including pyrimidine, purine, pyruvate, and citrate metabolism. (H) Comparison of basal metabolite abundance between TKI-sensitive and TKI-resistant cell lines, demonstrating increased abundance of multiple nucleotide metabolites in SLR cells. (I) Hierarchical clustering of significantly altered metabolites in SLR cells following VCB1 treatment, showing metabolic perturbations that overlap with those observed in SUPM2 cells but with lower magnitude. Statistical significance was defined as q<0.05.

### VCB1 produces extensive lipidomic remodeling distinct from BMS-303141

We next performed global lipidomic profiling of SUPM2 parental, and SLR cells treated with VCB1, BMS, or DMSO. Principal component analysis demonstrated clear treatment-dependent separation in both cell lines. In parental cells, DMSO, VCB1, and BMS samples formed distinct clusters, with VCB1 separating predominantly along PC1 and BMS along PC2 . A similar pattern was observed in SLR cells (Fig. 6A and B). In parental cells, VCB1 induced pronounced changes in multiple lipid species. The results showed that at using a stringent >15-fold-change threshold, the most prominent increased lipid species involved in sphingolipids, particularly ceramides and hexosylceramides, together with SM(d41:0), Hex2Cer(d18:0_24:0), LPE(16:0e), and PE(18:1p_20:4). Only three species PG(12:0_14:0), PG(10:0_14:0), and SM(d39:6) were strongly decreased (Fig. 6C–E). A similar pattern was observed in SLR cells, in which ceramides and related sphingolipids represented a major fraction of the most strongly increased species (Fig. 6F). These changes were substantially weaker following BMS treatment.

**Figure 6.**
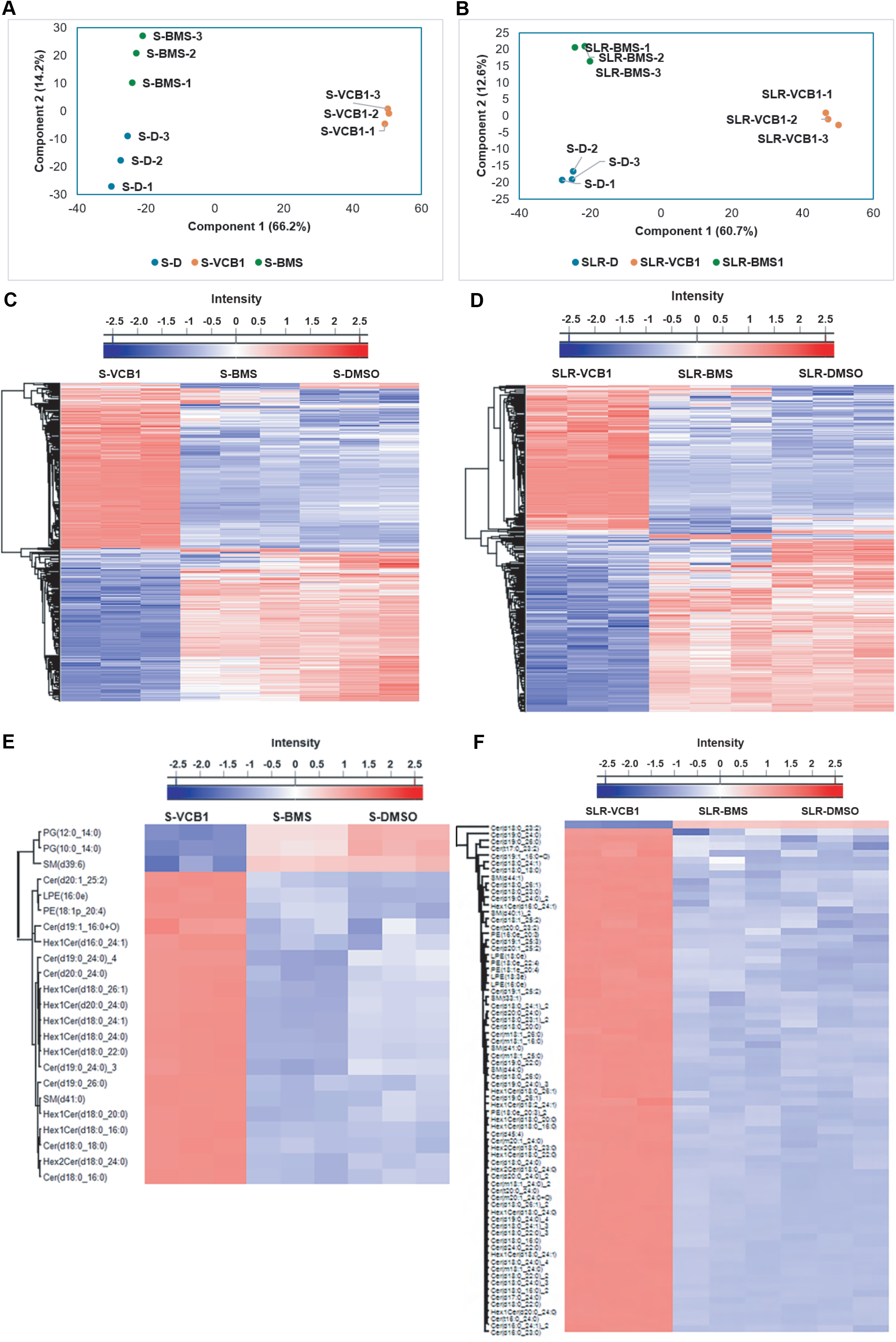
ACLY inhibition revealed distinct lipid remodeling in TKI-sensitive and TKI-resistant lymphoma cell lines. (A) PCA of untargeted lipidomic profiles from SUPM2 parental cells treated with DMSO, VCB1, or BMS-303141 (BMS), showing treatment-dependent separation of lipidomic profiles. (B) PCA of lipidomic profiles from SLR cells treated with DMSO, VCB1, or BMS. (C,D) Hierarchical clustering heat maps of significantly altered lipid species (q ≤ 0.05) in SUPM2 parental (C) and SLR (D) cells. (E) Heatmap of lipid species showing the most pronounced changes in SUPM2 parental cells following VCB1 treatment using a stringent >15-fold-change threshold. VCB1 treatment predominantly increased ceramide, hexosylceramide, sphingomyelin, and selected phospholipid species, whereas PG(12:0_14:0), PG(10:0_14:0), and SM(d39:6) were markedly decreased. (F) Heatmap of the most strongly altered lipid species in SLR cells using the same >15-fold-change threshold, demonstrating prominent increases in ceramide and related sphingolipid species following VCB1 treatment. Comparable changes were not observed in BMS or DMSO-treated cells. Each point in the PCA represents an individual biological replicate.

### ACLY Inhibition by VCB1 Reprograms Lipid Metabolism and Mitochondrial Fatty-Acid

The lipidomics analysis revealed that remodeling of lipid species across acylcarnitines, ceramides, glycosphingolipids, PC, PE, PG, cardiolipin, PS, PI, SM, DG, cholesteryl esters, and TG (Fig. 7A). In sensitive cells, numerous TG species containing oleate, very-long-chain fatty acids, and mixed unsaturated chains were depleted, whereas selected saturated C16–C18, ether-linked, and PUFA-containing TGs increased (Fig. 7B). PC species also shifted toward ether and PUFA-containing species, including PC(16:0e/20:4), PC(18:0e/22:6), and PC(16:1e/20:4), with depletion of conventional diacyl PCs (Fig. 7C). PE exhibited a particularly strong shift toward ether/plasmalogen- and PUFA-rich species, including PE(18:0e/20:4), PE(16:0e/20:5), PE(18:1p/20:4), PE(18:0e/22:5), and PE(18:0e/22:6) (Fig. 7D). DG species were extensively remodeled in both cell lines, whereas BMS produced comparatively weak changes (Fig. 7E). Further, ChE species were also broadly altered, with different responses between sensitive and resistant cells (Fig. 7F). VCB1 increased long-chain saturated acylcarnitines, including AcCa(17:0), (18:0), (19:0), (20:0), (22:0), and (24:0), while decreasing shorter-chain and unsaturated species including AcCa(9:0), (10:0), (12:1), (14:0), (16:1), (18:1), and (18:2) (Fig. 7G), consistent with altered mitochondrial fatty-acid handling. Mitochondrial and membrane phospholipids were also remodeled, including predominant depletion of identified cardiolipin and PS species. Acylcarnitines are intermediates required to transport fatty acids into the mitochondria for energy production. The accumulation of very-long and long-chain species, paired with the drop in shorter-chain species, indicates a "bottleneck" in mitochondrial beta-oxidation. VCB1 likely blocks or overwhelms the enzymes responsible for breaking down long-chain fatty acids, leading to mitochondrial congestion and severe bioenergetic stress. Collectively, these results establish that VCB1 produces substantially broader lipid remodeling than BMS, encompassing neutral lipids, membrane phospholipids, mitochondrial lipids, acylcarnitines, sphingolipids, and cholesterol-associated species.

**Figure 7.**
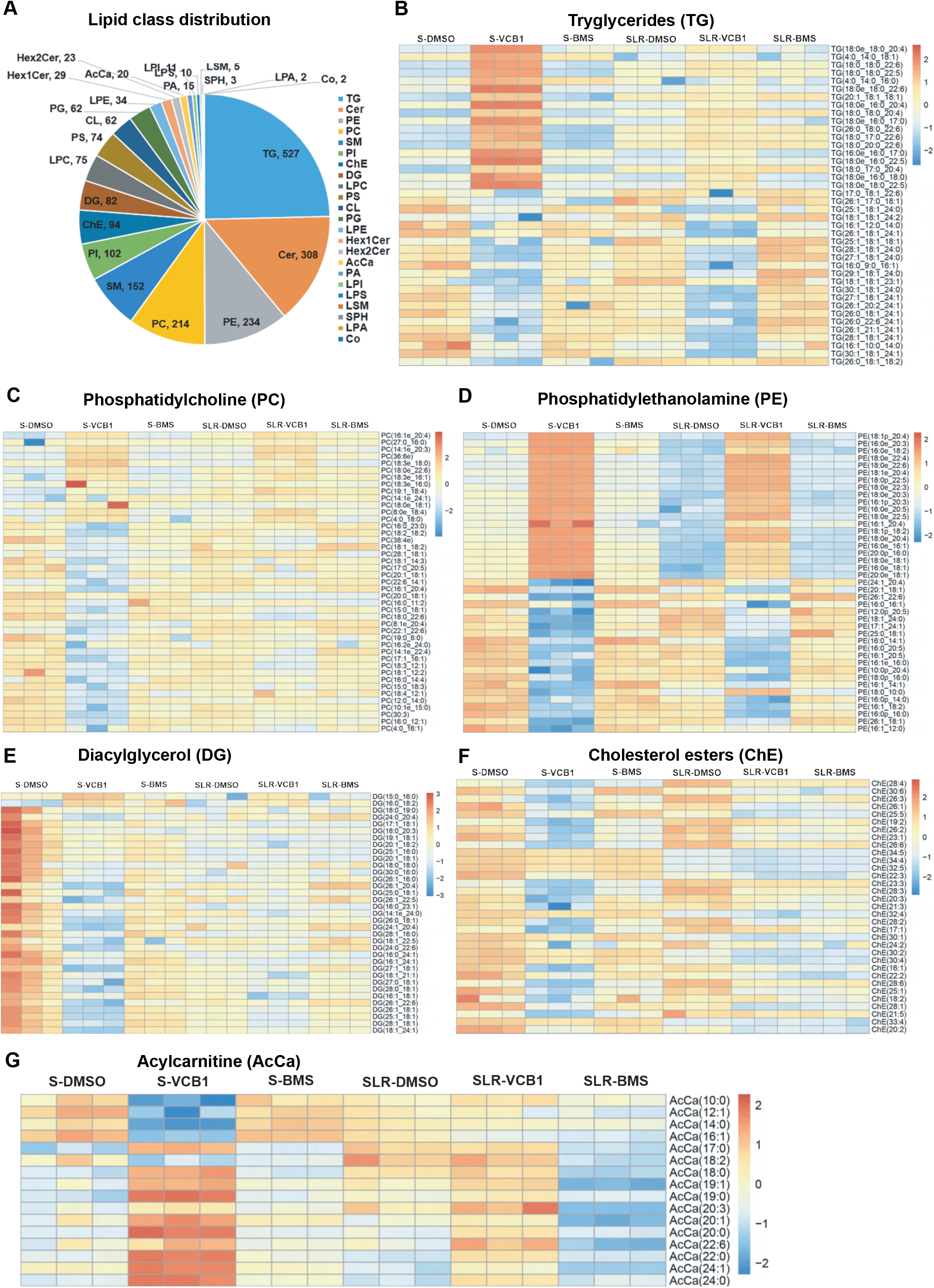
VCB1-mediated ACLY inhibition induces extensive lipid species and transcriptional remodeling in lymphoma cells. (A) Distribution of lipid classes identified by untargeted lipidomic profiling following ACLY inhibition. VCB1 induced broad remodeling across acylcarnitines, ceramides, glycosphingolipids, PC, PE, PG, CL, PS, PI, SM, DG, ChE, and TG classes. (B) Differential abundance of TG molecular species, demonstrating more pronounced remodeling in SUPM2 cells, with reciprocal changes in species containing saturated, unsaturated, very-long-chain, ether-linked, and PUFA-containing fatty acids. (C) Remodeling of PC species, characterized by enrichment of selected ether-linked and PUFA-containing PCs and depletion of multiple conventional diacyl PCs. (D) Remodeling of PE species, with preferential enrichment of ether/plasmalogen and PUFA-containing species and depletion of selected shorter-chain saturated/monounsaturated species. (E) Differential abundance of DG species following VCB1 treatment in SUPM2 and SLR cells. VCB1 produced broad, species-specific DG remodeling, whereas BMS induced comparatively weak changes. (F) Differential abundance of ChE species following VCB1 or BMS treatment, demonstrating broad remodeling of cholesterol ester species following VCB1 treatment. (G) Changes in acylcarnitine species, showing accumulation of several long-chain saturated species and depletion of shorter-chain and unsaturated species, consistent with altered mitochondrial fatty-acid handling.

### ACLY inhibition induces metabolic remodeling, integrated stress response, redox stress, apoptosis, and immune regulation

To define the transcriptional consequences of VCB1-mediated ACLY inhibition, DEGs associated with lipid metabolism were identified using a log2 FC ≥0.5 threshold and NIH-DAVID/KEGG pathway analysis. VCB1, but not BMS-303141 at the same concentration, significantly altered lipid-metabolic pathways and enzymes (Fig. 8A). These findings prompted analysis of the integrated stress response (ISR), which couples translational control to metabolic and amino-acid adaptation through eIF2α kinase signaling(47–51). VCB1 induced a prominent ISR-associated transcriptional response in both ALKi-sensitive and -resistant cells, including *ATF4, ATF3, SESN2, DDIT4, HILPDA, TRIB3, DDIT3, PPP1R15A,* and *CHAC1* (log2 FC +2.3 and +1.8 in S-VCB1 and SLR-VCB1), whereas ASNS was additionally induced only in sensitive cells (log2 FC = 1.25) and *BNIP3* (log2 FC +2.0 and +1.6) and *BNIP3L* (log2 FC +1.6 and +1.7) indicated mitochondrial stress adaptation (Fig. 8B). VCB1 induced *BNIP3L* mRNA expression results showed impaired lipid synthesis. Similarly, in another study confirmed *BNIP3L* protein, was significantly reduced in patients with Diabetic cardiomyopathy compared to those with non-diabetic(52). BMS-303141 produced no significant ISR-associated transcriptional response at 1 μM, although activation of the p-eIF2α/ATF4/CHOP pathway has been reported at higher concentrations in hepatocellular carcinoma(53). Furthermore, our results demonstrated a conserved ISR with distinct downstream stress-adaptation programs in SUPM2 and SLR cells. RT-PCR confirmed VCB1-induced changes in *PPP1R15A/GADD34, NLRP3,* and *CHAC1* observed by RNA-seq data (Fig. 8C-E). *PPP1R15A/GADD34* is a key regulator of ISR termination(54), whereas CHAC1 promotes glutathione degradation and contributes to oxidative stress and cell death(55). Consistent with this response, VCB1 increased CHAC1 by +2.26 and +1.77 log2 FC in sensitive and resistant cells (Fig. 8E). Metabolomics analysis demonstrated corresponding disruption of glutathione homeostasis, with reduced GSH and GSSG and a significantly decreased GSH/GSSG ratio following VCB1 treatment, whereas BMS-303141 produced no significant change (Fig. 8F). VCB1 also decreased γ-glutamylcysteine and S-lactoylglutathione (Fig. 8G-H). Thus, VCB1 treatment showed concordant transcriptional and metabolic evidence of impaired glutathione homeostasis, supporting an ACLY–ISR–CHAC1 axis linking metabolic stress to redox imbalance. Consistent with activation of cell-death pathways, VCB1 also increased apoptosis/stress-associated genes including *ATF4, GADD45A, GADD45B, DDIT3, DDIT4, GZMA, TRAF1,* and *TNF* (Fig. 8I). VCB1 also induced inflammatory and immune-regulatory remodeling. *NLRC5* was reduced, whereas *NLRP1* and *NLRP3* were increased in both sensitive and resistant cells, with strong *NLRP3* induction in the resistant state (Fig. 8J). The activating immune receptor *CD226* increased in sensitive (+1.49 log2 FC) and resistant (+0.67 log2 FC) cells, while *PDCD1* (PD-1) particularly increased in resistant cells (+1.19 log2 FC). *PDCD4* was also induced (+0.89 log2 FC), accompanied by increased *EIF4EBP1* and reduced *EIF4A1*, consistent with suppression of eIF4A-dependent translation during metabolic stress. Although PD-L1 was not differentially expressed in our RNA-seq dataset, previous studies reported increased PD-L1 following ACLY inhibition by bempedoic acid or genetic ACLY knockdown(56). Together, these findings indicate that ACLY inhibition alters inflammatory, immune-checkpoint, and translational regulatory pathways. The relationship between *PDCD1* and *NLRP3* involved in ISR, and ER stress further supports a link between metabolic stress and immune regulation(57). Finally, integration of VCB1-responsive RNA-seq and metabolomics data using MetaboAnalyst 6.0 identified significant gene-metabolite interactions involving lipid metabolic pathways in sensitive cells (Fig. 8K), with a corresponding network observed in resistant cells (Supplementary Fig. 7). The integrated model summarizes VCB1-mediated ACLY inhibition as a coordinated response involving altered central-carbon metabolism, glycolytic and amino-acid metabolic genes (*PFKFB3/4, ENO2, ASNS*), suppression of ACLY-dependent lipogenesis (*ACACA, FASN*), and lipid-remodeling pathways (*HILPDA, DGKG*), accompanied by lipid-species remodeling, ISR activation, glutathione/redox stress, apoptosis, and inflammatory and T-cell regulatory signaling (Fig. 8L).

**Figure 8.**
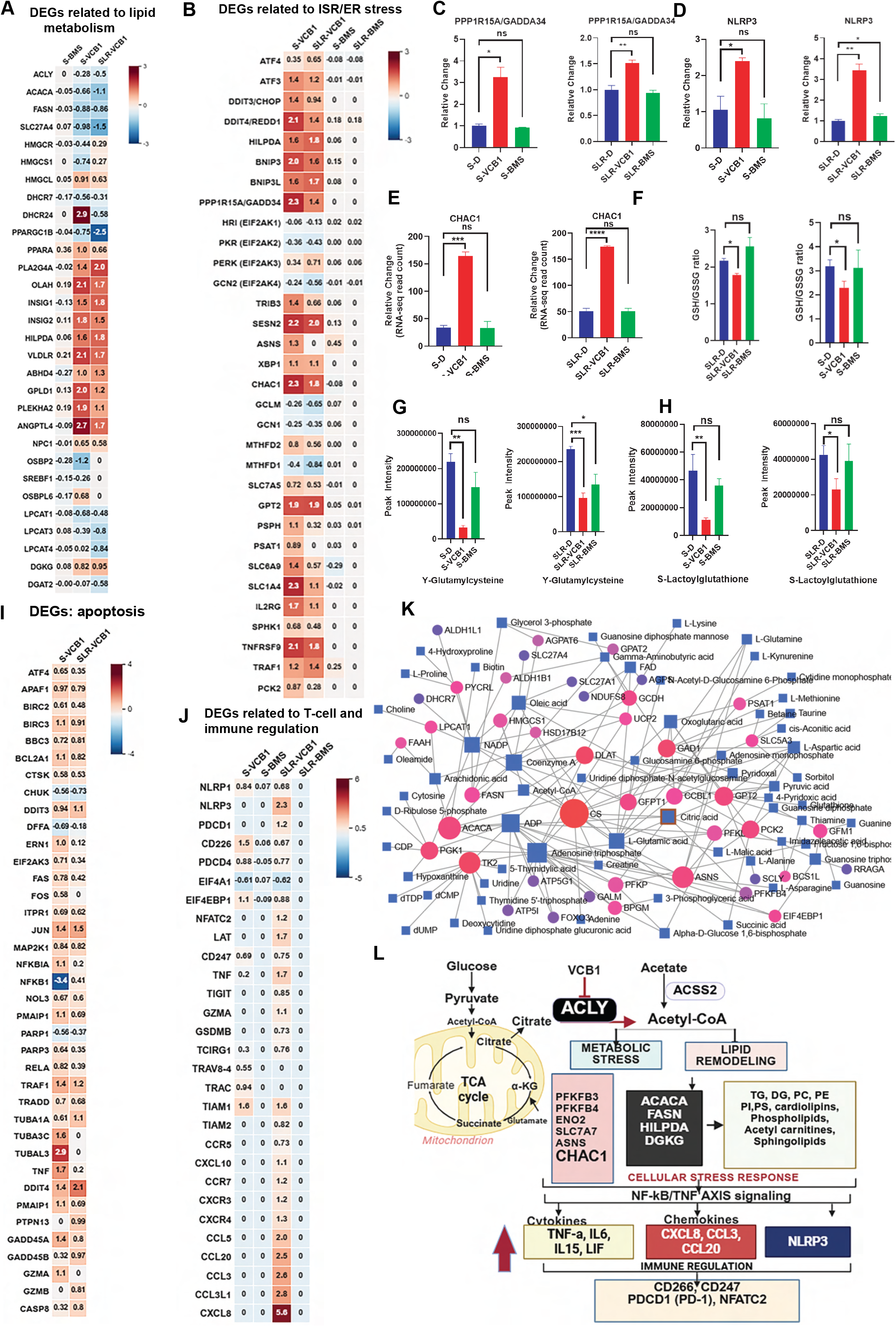
VCB1 induces metabolic remodeling, integrated stress response, redox stress, apoptosis, and immune regulation. (A) Heatmap based on the NIH-DAVID and KEGG pathway analysis of lipid-metabolism-associated DEGs following VCB1 treatment, demonstrating significant remodeling of lipid metabolic pathways that were not observed with BMS-303141 at the same concentration. (B) Heatmap of representative ISR/stress-associated DEGs induced by VCB1 (C-E) RT-PCR validation of representative ISR, inflammatory, and redox-associated genes, including PPP1R15A/GADD34, NLRP3, and CHAC1, respectively. (F-H) Effects of VCB1 and BMS-303141 on glutathione metabolism, including GSH/GSSG ratio, γ-glutamylcysteine, and S-lactoylglutathione levels. (I) Representative apoptosis- and stress-associated DEGs induced by VCB1. (J) Differential expressions of inflammatory and immune-regulatory genes, including NLRC5, NLRP1, NLRP3, CD226, PDCD1, and PDCD4. (K) Integrated gene-metabolite interaction network generated from VCB1-responsive transcriptomic and metabolomic datasets demonstrating enrichment of lipid-metabolic interactions in sensitive cells. (L) Schematic model summarizing the downstream consequences of VCB1-mediated ACLY inhibition, including metabolic remodeling, suppression of ACLY-dependent lipogenesis, ISR activation, CHAC1-associated glutathione depletion, redox and apoptotic stress, and inflammatory/immune-regulatory signaling. Unless otherwise indicated, differential expression was defined using a log2 FC threshold of ≥0.5 or ≤−0.5.

### VCB1 shows potent antiproliferative activity in tyrosine kinase inhibitor-resistant lymphoma

We next evaluated the antiproliferative activity of VCB1, VCB2, VCB7, and BMS-303141 in SUPM2 parental and lorlatinib-resistant SUPM2-LR1000 cells. Cells were treated with increasing concentrations of each compound for 72 h in 5% FBS. VCB1 showed the strongest growth-inhibitory activity in SUPM2 parental cells, with an IC50 of 0.299 μM, compared with 22.0 μM for VCB2, 1.21 μM for VCB7, and 17.46 μM for BMS. In SLR cells, VCB1 retained substantial activity, although its potency was reduced, with an IC50 of 1.62 μM. VCB2, VCB7, and BMS showed IC50 values of 9.98, 3.60, and 22.07 μM, respectively (Fig. 9A, B). VCB1 also displayed potent activity in MCL models. In Jeko-1 and Maver cells, VCB1 showed IC50 values of 0.28 and 0.278 μM, respectively. In contrast, VCB2 showed IC50 values of 15.0 and 10.4 μM, VCB7 showed values of 2.26 and 1.84 μM, and BMS showed substantially weaker activity, with IC50 values of 30.5 and 37.01 μM, respectively (Fig. 9C, D). Thus, VCB1 was consistently the most potent compound tested and retained activity in lymphoma models with acquired resistance to ALK or BTK inhibition. These findings indicate that ACLY inhibition represents a therapeutically distinct vulnerability that can be maintained despite resistance to kinase-targeted therapy. Next, we investigated the RNA-seq data to determine whether the transcriptional and metabolic alterations translated into functional effects on proliferation, we examined cell-cycle progression related DEGs in cells treated with VCB1. The results showed significant changes in both ALKi sensitive and resistant cell lines (Fig.9E). Further, we performed cell cycle analysis on these cell lines treated with 0.5 μM VCB1 for 48 h resulted in significant accumulation of cells in the G0/G1 phase compared with BMS-treated cells (Fig. 9F–H). These findings are consistent with the RNA-seq suppression of CCNB1, CDC20, PLK1, CENPE, CENPF, KIF20A, KIF23, and PTTG1 and induction of cell-cycle checkpoint genes such as CDKN1A, GADD45A, and BTG2. Because metabolomics revealed substantial disruption of pyrimidine metabolism, we examined key enzymes within this pathway. RNA-seq demonstrated reduced expression of DHODH and CAD following VCB1 treatment, (Fig. 9I, J). RNA-seq further demonstrated coordinated disruption of pyrimidine biosynthesis DEGs following VCB1, whereas BMS produced substantially weaker changes (Fig. 9K). Further analysis on RNA-seq identified DEGs involved in phospholipid, LPCAT1 (Fig.9L), cholesterol synthesis pathway DEGs, HMGCR, INSIG1 and INSIG2 (Fig.9L-O ). In summary, Integration of RNA-seq and metabolomic datasets further demonstrated coordinated disruption of pyrimidine biosynthesis following VCB1, whereas BMS produced substantially weaker changes (Fig. 9P). Together, these findings indicate that VCB1-mediated ACLY inhibition disrupts interconnected lipid, nucleotide, central-carbon, redox, and stress pathways, resulting in cell-cycle arrest and ultimately apoptotic cell death.

**Figure 9.**
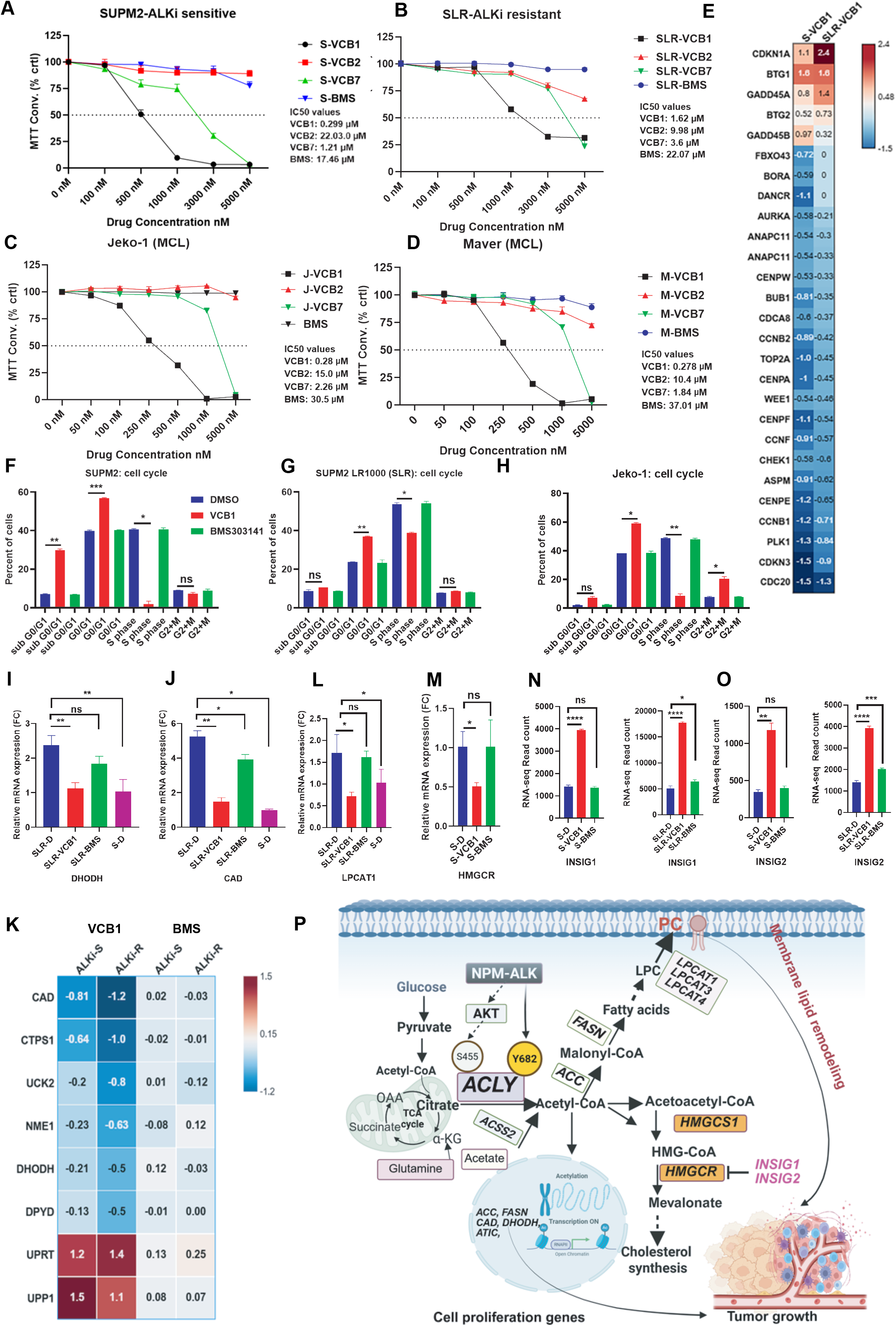
VCB1 inhibits lymphoma cell proliferation through coordinated metabolic stress induced response and cell-cycle alteration. (A,B) Dose–response analysis of VCB1, VCB2, VCB7, and BMS in TKI-sensitive, SUPM2 and TKI-resistant, SLR lymphoma cells following 72 h treatment in 5% FBS. (C,D) Antiproliferative activity of the indicated ACLY inhibitors in BTK inhibitor-sensitive Jeko-1 and BTK inhibitor-resistant Maver mantle cell lymphoma cells. (E) RNA-seq analysis of differentially expressed genes associated with cell-cycle progression and cell growth following VCB1 treatment. (F–H) Cell-cycle analysis of SUPM2 parental, SLR, and Jeko-1 cells treated with 0.5 μM VCB1 or BMS for 48 h, demonstrating increased G0/G1-phase accumulation following VCB1 treatment. (I, J) RNA-seq and RT-qPCR validation of DHODH and CAD expression following VCB1 treatment. (K) Differential expression of pyrimidine biosynthesis-related genes following VCB1 treatment compared with BMS-303141. (L–O) Expression of LPCAT1 and cholesterol biosynthesis-associated genes, including HMGCR, INSIG1, and INSIG2, following VCB1 treatment. (P) Integrated analysis of RNA-seq and metabolomics demonstrating coordinated disruption of pyrimidine biosynthesis by VCB1. Data are presented as mean ± SEM; statistical significance was determined using the graph prism software and performed t-test analysis. A p-value of < 0.05 was considered statistically significant (*, p <0.05; **, p < 0.01; ***, p < 0.001; ****, p < 0.0001).

**Table 1.** Common DEGs in TKI sensitive and resistant cell lines. Common metabolic function related differentially expressed genes (DEGs) in TKI-sensitive, SUPM2 and TKI-resistant, SLR cells following VCB1 treatment. Metabolic function-associated differentially expressed genes (DEGs) were identified from RNA-seq data using a log2 fold-change (FC) threshold of ≥ +0.5 or ≤ −0.5 in both cells following VCB1 treatment.

| Gene symbol | S-VCB1<br>(log<sub>2</sub> FC) | SLR-VCB1<br>(log<sub>2</sub> FC) | Metabolic function |
| --- | --- | --- | --- |
| PFKFB4 | 2.73 | 2 | Glycolysis; regulates fructose-2,6-bisphosphate and metabolic stress adaptation |
| PFKL | 0.89 | 0.9 | Glycolysis; rate-limiting phosphofructokinase activity |
| GPAT4 | 0.52 | 0.84 | Glycerolipid/triglyceride synthesis |
| GAD1 | 1.15 | 1.25 | Glutamate → GABA metabolism |
| PPARGC1B | -0.751 | -2.497 | PGC-1β; mitochondrial oxidative metabolism, fatty-acid oxidation and energy metabolism |
| ABCD2 | -1.388 | -1.239 | Peroxisomal fatty-acid transport and β-oxidation |
| ALDH1B1 | -0.683 | -1.237 | Aldehyde metabolism; mitochondrial metabolic/detoxification function |
| **ACACA** | -0.660 | -1.149 | Acetyl-CoA → malonyl-CoA; rate-limiting de novo fatty-acid synthesis |
| SCLY | -0.953 | -1.128 | Selenocysteine/selenium recycling; antioxidant metabolism |
| BCS1L | -0.586 | -1.121 | Mitochondrial complex III assembly and oxidative phosphorylation |
| **FASN** | -0.884 | -0.859 | De novo fatty-acid synthesis; converts acetyl-CoA/malonyl-CoA into fatty acids |
| CS | -0.543 | -0.739 | TCA cycle; citrate production from acetyl-CoA and oxaloacetate |
| BPGM | -0.906 | -0.802 | Glycolytic intermediate/2,3-BPG metabolism |
| AGPS | -0.506 | -0.642 | Ether phospholipid/plasmalogen biosynthesis |
| HACD2 | -0.525 | -0.597 | Fatty-acid elongation; very-long-chain fatty-acid synthesis |
| PYCR3 | -0.599 | -0.539 | Proline biosynthesis; NAD(P)H-dependent reduction of P5C |
| DLAT | -0.503 | -0.599 | Pyruvate dehydrogenase complex; pyruvate → acetyl-CoA |
| NDUFS8 | -0.514 | -0.508 | Mitochondrial complex I/electron transport |
| COQ2 | -0.563 | -0.631 | Coenzyme Q biosynthesis; mitochondrial respiration |

## Discussion

ACLY occupies a central metabolic position by converting citrate to cytosolic acetyl-CoA, thereby linking glucose-derived carbon to fatty-acid and cholesterol synthesis. However, therapeutic targeting of ACLY remains challenging. Bempedoic acid (also known as ETC-1002) has been recently approved by U.S. Food and Drug Administration (FDA) for lowering low-density lipoprotein cholesterol(27, 58). Bempedoic acid targeting ACLY-requires activation by SLC27A2/ACSVL1, is predominantly active in the liver, and activates AMPK pathway(59, 60). Like this other ACLY inhibitor, EVT0185 shows anticancer activity but requires metabolic activation and has additional activity against ACSS2(60, 61). In our studies, the reference ACLY inhibitor BMS-303141 inhibited recombinant ACLY but showed limited cellular activity in ALCL and MCL models. These limitations highlight the need for potent, selective, cell-active ACLY inhibitors.

Building on our previous demonstration that NPM-ALK phosphorylates ACLY at Y682 to promote lipid metabolism and ALCL progression(6, 30, 31) and that ACLY is regulated downstream of the PI3K/PIP2/PIP3–LYN axis(32), we identified VCB1 as a novel, cell-active ACLY inhibitor. Our xenograft findings provide genetic *in-vivo* evidence supporting the role of ACLY in ALK-positive ALCL tumor establishment. Although ACLY is an established regulator of cancer metabolism and tumor growth, genetic validation of ACLY dependence in lymphoma xenograft models remains limited. In the SU-DHL-1 model, control cells eventually established substantial tumors, whereas ACLY-depleted cells failed to produce detectable tumors during the observation period. Thus, ACLY depletion was associated with a marked impairment of tumor establishment rather than simply a modest reduction in tumor growth and provide a rationale for pharmacologic targeting of ACLY in ALK-positive ALCL. We subsequently evaluated our novel ACLY inhibitor VCB1 in the paired SUP-M2 parental and lorlatinib-resistant SUP-M2-LR1000 models, which provided a common cellular background for investigating ACLY inhibition across ALK-inhibitor-sensitive and resistant states and for subsequent transcriptomic, metabolomic, and lipidomic analyses. VCB1 inhibited recombinant ACLY with an IC₅₀ of ∼0.9 µM and exhibited potent antiproliferative activity in ALK+ lymphoma, including TKI-resistant ALCL and BTK inhibitor-resistant MCL models. In SUPM2 cells, VCB1 showed a growth IC₅₀ of 0.299 μM and induced apoptosis, whereas BMS-303141 displayed substantially weaker cellular activity. Notably, BMS-303141 has generally been evaluated at relatively high micromolar concentrations in published cellular studies, including 10 μM in B-ALL(62), 5–15 μM in T-cell lymphoma(63), 10 μM in T-ALL(18), breast cancer(64), prostate cancer(12), and 25–50 μM in osteosarcoma models(53, 65). Similarly, the classical ACLY inhibitor SB-204990(22, 66) has generally required 10–100 μM concentrations in cellular studies, including 30 μM in HepG2 cells, 25 μM in human macrophages, 10–30 μM in ovarian cancer, 10–20 μM in lung cancer, 40 μM in KRAS-driven HCT116 cells, and 100 μM in pancreatic β-cells and other models(67). Although differences in permeability, intracellular exposure, and experimental systems preclude direct potency comparisons, VCB1 produced a growth IC₅₀ of 0.299 μM in SUPM2 cells and robust apoptotic signaling at 1 μM, supporting its characterization as a highly cell-active ACLY inhibitor and providing a rationale for further development in TKI-resistant lymphoma.

VCB1-mediated ACLY inhibition extended beyond de novo lipogenesis. Suppression of ACACA, FASN, HMGCR, and PPARGC1B, together with changes in SLC27A4 and CPT1A, was accompanied by induction of metabolic-adaptation genes including PPARA, ACADS, ACADVL, HMGCL, LPL, LDLR, VLDLR, and OSBPL5. PPARGC1B, encoding PGC-1β, regulates lipid and energy metabolism through SREBP and ChREBP(68) and has been implicated in glycolytic regulation and tumor-supportive macrophage phenotypes(69). PPARGC1B stronger downregulation in resistant cells (−2.5 versus −0.75 log2 FC in sensitive cells) is consistent with impaired mitochondrial and oxidative metabolic programs, similar observations in ACLY knockdown conditions(70). Hallmark analysis further identified activation of glycolytic and metabolic-stress programs, including SLC2A1, SLC2A3, PDK1, PGK1, PFKFB4, CA9, DDIT4, BNIP3L, and HILPDA. Together with changes in acetyl-CoA, citrate, acetylcarnitines, and TCA intermediates, these findings indicate that ACLY inhibition induces broad metabolic rewiring rather than simply suppressing fatty-acid synthesis.

Integrated transcriptomic, metabolomic, and lipidomic analyses revealed that SUPM2 and SLR lymphoma cells share a core response to ACLY inhibition but engage distinct metabolic adaptations. Thus, SLR appears to represent metabolic rewiring that changes how cells respond to ACLY blockade rather than simply a generalized increase in metabolism. In SUPM2 cells, VCB1 predominantly induced an acute biosynthetic and metabolic-stress response involving glycolysis, lipid metabolism, amino-acid metabolism, and mTOR/AMPK signaling. Suppression of ACACA, FASN, HMGCR, CS, DLAT, and PPARGC1B together with induction of PFKFB4 and PFKL suggests increased reliance on alternative energy-generating pathways following loss of ACLY-derived acetyl-CoA. In contrast, SLR cells exhibited more extensive metabolic remodeling, including increased expression of genes associated with glucose and glutamine utilization, lipid uptake and storage, cholesterol metabolism, and fatty-acid utilization, including LDLR, APOC1, DGAT2, FADS2, and DHCR24, while mitochondrial, TCA-cycle, fatty-acid, and nucleotide metabolic programs were broadly suppressed. Importantly, sensitive and resistant cells shared induction of the integrated stress response (ISR), including ATF4, ATF3, DDIT3/CHOP, DDIT4, SESN2, TRIB3, PPP1R15A, and CHAC1. Thus, despite distinct basal metabolic states, both populations recognize effective ACLY inhibition as metabolic stress and activate a conserved ISR. This is consistent with the established role of ATF4 as a central ISR effector coordinating amino-acid, redox, and metabolic adaptation and inducing CHAC1-dependent glutathione depletion. In previous studies by others showed that ATF4 also regulates nucleotide synthesis and DNA replication in leukemia cells(71).

The conserved ISR produced distinct downstream adaptations. Sensitive cells showed greater acute ER/UPR and metabolic-stress responses, whereas resistant cells exhibited stronger PERK–ATF4, glutathione/redox, autophagy, and apoptotic programs. This suggests that the consequences of ACLY inhibition are shaped by pre-existing metabolic state. In resistant cells, extensive metabolic flexibility may permit survival following kinase inhibition but simultaneously limit compensation for disruption of lipid synthesis, mitochondrial metabolism, and redox homeostasis. Lipidomics analysis further demonstrated that VCB1 does not simply suppress lipid production but reorganizes lipid metabolism and membrane composition. VCB1 altered triglycerides, phospholipids, cholesteryl esters, sphingolipids, cardiolipins, and acylcarnitines, with changes in DGK, GPAT, LPCAT, PLD, CHKA/E, PTDSS1, CDS, and DGAT2 indicating remodeling across multiple lipid pathways. Ceramide and hexosylceramide accumulation in both cellular states, together with cardiolipin and phosphatidylglycerol alterations and increased acylcarnitines, suggests disruption of mitochondrial lipid homeostasis. These findings extend previous observations linking ACLY inhibition to mitochondrial dysfunction, including impaired membrane potential and oxidative phosphorylation(21, 72). A particularly important, shared consequence was disruption of glutathione homeostasis. CHAC1 was strongly induced in sensitive (+2.26 log2 FC) and resistant (+1.77 log2 FC) cells, accompanied by reduced γ-glutamylcysteine, glutathione, and GSH/GSSG ratio. Because ATF4 can induce CHAC1 and promote GSH degradation, these findings provide a mechanistic link between the ISR and redox stress, supporting an ACLY–ISR–CHAC1–redox axis as a potential mechanism underlying oxidative vulnerability following ACLY inhibition. VCB1 also disrupted nucleotide metabolism. Metabolomics revealed depletion or alteration of AMP, dAMP, dADP, UMP, TMP, cytosine, GMP, IMP, GTP, CMP, and CTP, accompanied by transcriptional changes in CAD, DHODH, PPAT, GMPS, ADSL, ATIC, CTPS, and UMPS. Reduced aspartate and altered orotic acid and orotidine further support impaired de novo pyrimidine synthesis. The convergence of transcriptional and metabolite changes indicates that ACLY inhibition disrupts nucleotide supply in addition to lipid synthesis, potentially limiting the biosynthetic capacity of rapidly proliferating lymphoma cells. Consistent with this, VCB1 induced G0/G1 accumulation and increased CDKN1A, GADD45A, and BTG2 while suppressing proliferation-associated genes. Thus, VCB1-mediated growth inhibition likely reflects coordinated disruption of lipid, nucleotide, and redox homeostasis rather than lipogenesis blockade alone. Lipid metabolism also influences immune-cell function and innate and adaptive immune responses(73). In this context, VCB1 induced inflammatory and stress-associated genes in both cell lines, including increased NLRP1 and NLRP3 and reduced NLRC5, with particularly strong NLRP3 induction in resistant cells (+2.50 log2 FC). Because NLRP3 responds to metabolic and cellular stress, its greater induction may reflect the increased metabolic burden of resistant cells, although its contribution to VCB1-mediated cytotoxicity remains to be established. Changes in PDCD4, CD226, and PDCD1 (PD-1), together with altered EIF4-associated factors, further indicate translational and immune-associated remodeling. The results showed that VCB1 induced PD-1 expression in SLR cells (+1.2 log2 FC). The previous study showed that the association of PD-1 signaling suppressing ACLY-acetyl-CoA production and acts as tumor suppressor(63). Similarly, VCB1 treatment consistently induced PDCD4 level (+0.89 log2 FC), accompanied by increased EIF4EBP1 and reduced EIF4A1, supporting suppression of protein synthesis during metabolic stress. PDCD4 is a recognized tumor suppressor involved in translational control, apoptosis, and inflammatory signaling(74, 75), and its loss promotes B-cell lymphoma development in mice(76). Translational repression may therefore reinforce the ISR-associated reduction in anabolic growth. These findings support a model in which ACLY inhibition produces a conserved core stress response superimposed on distinct metabolic programs determined by TKI sensitivity or resistance. Common VCB1-responsive DEGs reflect suppression of anabolic metabolism, activation of the ATF4/ATF3/CHOP ISR, CHAC1-associated glutathione depletion, inflammatory signaling, and translational repression. In contrast, sensitivity-specific responses reflect acute biosynthetic and metabolic stress, whereas resistance-specific responses reveal compensatory dependence on glycolysis, glutamine utilization, exogenous lipid acquisition, fatty-acid utilization, and redox metabolism. Thus, TKI resistance appears to be remodeled rather than simply increasing cellular metabolism. This rewiring may permit survival during kinase inhibition while constraining the capacity of resistant lymphoma cells to compensate for ACLY blockade. VCB1 therefore targets a metabolic node that remains important despite loss of kinase dependence, while resistance-associated adaptations may create additional vulnerabilities. The shared ISR/redox stress together with resistance-specific metabolic dependence provides a mechanistic framework for ACLY inhibition as a resistance-bypass strategy in lymphoma.

## Conclusion

Collectively, our findings extend the biological consequences of ACLY inhibition beyond lipid synthesis. We identify VCB1 as a novel, cell-active ACLY inhibitor with sub micromolar antiproliferative activity in TKI-sensitive lymphoma cells (IC₅₀ = 0.299 μM) and retained activity against lorlatinib-resistant cells (IC₅₀ 1.62 μM), whereas BMS-303141 showed limited cellular activity in our models. Integrated transcriptomic, metabolomic, and lipidomic analyses demonstrate that VCB1 induces broad metabolic remodeling involving lipid and nucleotide metabolism, mitochondrial homeostasis, and redox balance. VCB1 also activates a robust ISR characterized by ATF4-associated stress genes, including CHAC1, ATF3, DDIT3, DDIT4, SESN2, and TRIB3. Notably, the stronger transcriptional response in TKI-resistant cells suggests that resistance-associated metabolic rewiring may increase dependence on ACLY-linked metabolic programs. Together, these findings define an ACLY–nucleotide metabolism–ISR axis connecting metabolic disruption with impaired biosynthetic and redox homeostasis and support ACLY inhibition, particularly with cell-active compounds such as VCB1, as a potential therapeutic strategy for TKI-resistant lymphoma.

## Acknowledgments

We thank the Proteomics and Metabolomics Shared Resource at the Wistar Institute, Philadelphia, for providing metabolomics and proteomics analysis services and for their support. The figures in this manuscript were created using the Biorender program. L.M., is supported by Italian Association for Cancer Research grant n. 24828 funding. H.Y.T is supported by R50CA221838. Funding support for The Wistar Institute’s Proteomics and Metabolomics Shared Resource was provided by Cancer Center Support Grant P30 CA010815. No NIH or other US federal funds were used to support co-authors with foreign affiliations.

## Author Contributions

J.B. conceived the project, designed and performed the research, analyzed the data, supervised the project, and wrote the manuscript. N.P., in-silico screening of ACLY inhibitors, performed molecular docking. C.L., D.R., and S.W., investigation, A.R.G. and W.Z., conducted metabolomics and lipidomics and data analysis. L.M., H.B., K.N. contributed to xenograft study, R.N. contributed to the resources and M.W. obtained funding and provided resources. All authors have approved the manuscript and agree with its submission.

## Supplementary Figure legends

**Supplementary Fig. 1.**
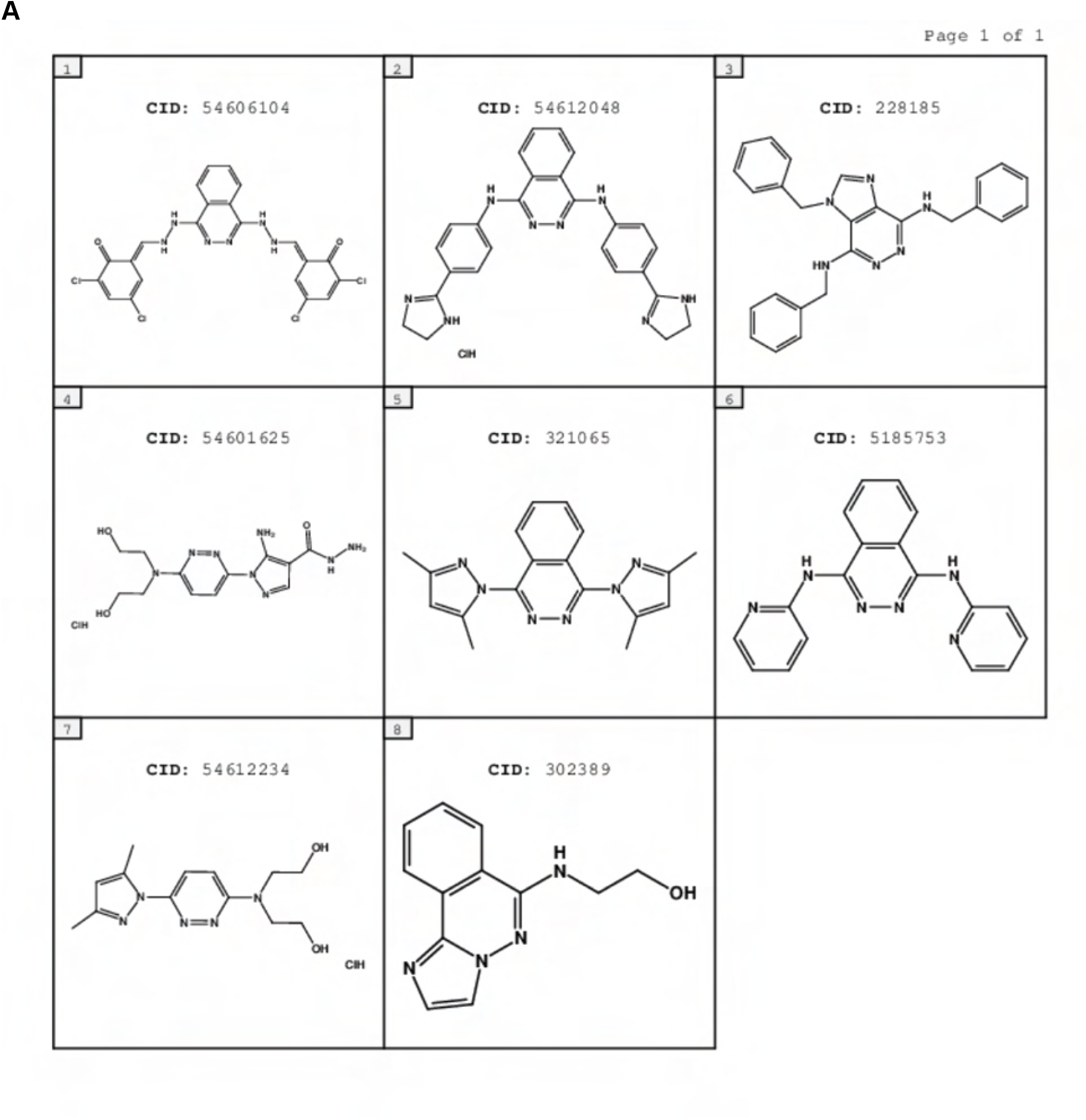
Chemical structures of ACLY inhibitor candidates selected for activity screening. Two-dimensional chemical structures of the additional compounds selected from the NCI repository for ACLY activity screening.

**Supplementary Fig. 2.**
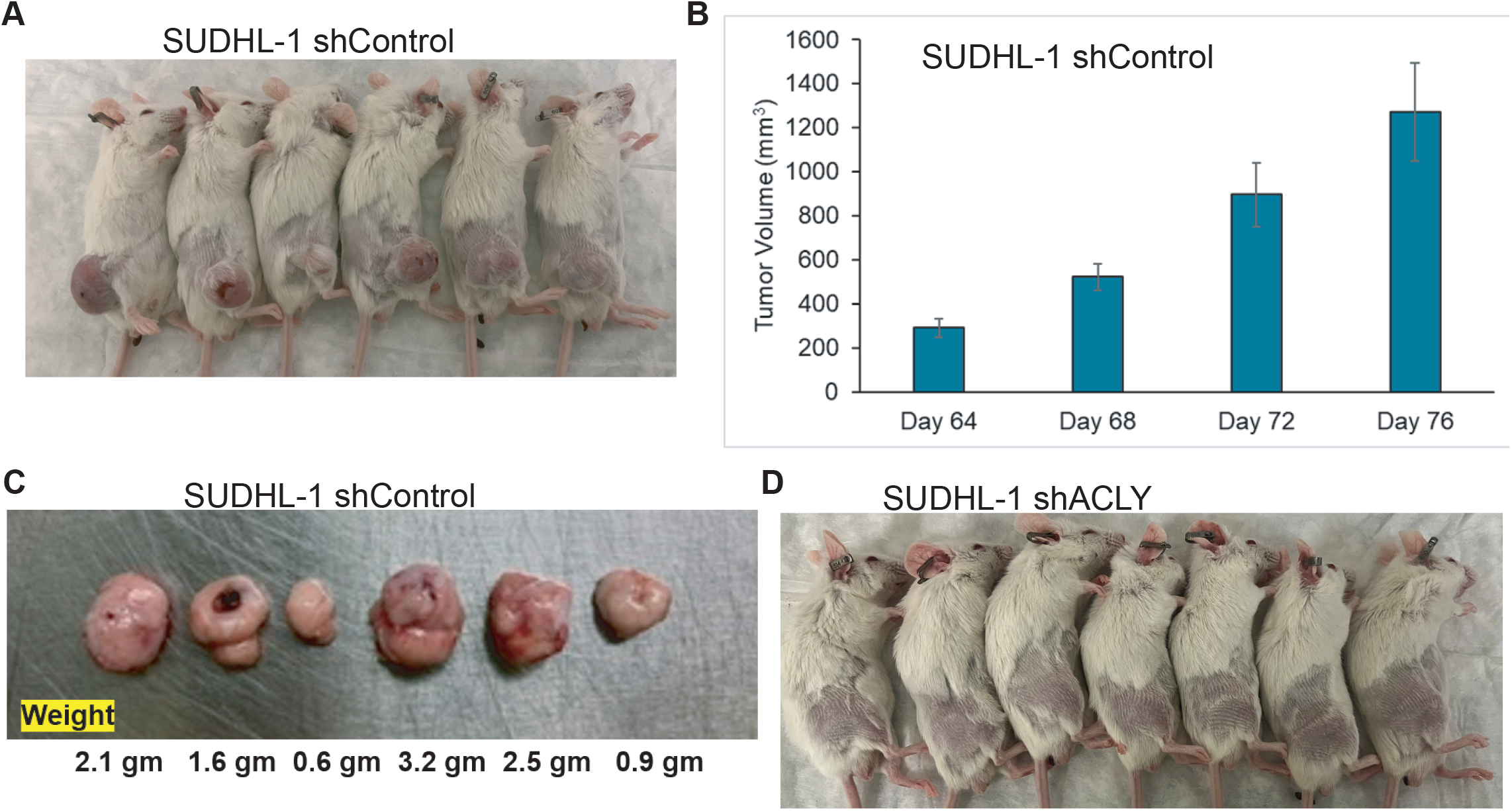
Genetic silencing of ACLY in ALK+ALCL cell line and xenograft model revealed ACLY as a therapeutic target (A) Images of tumor bearing mice derived from shControl SUDHL-1 cells 2X10^6^ cells xenografted into SCID-beige mice. (B) Tumor volumes of xenografts derived from representative images in figure A. (C) Representative tumor weights at the endpoint of the experiment. (D) Images of non-tumor bearing mice derived from shACLY SUDHL-1 cells 2X10^6^ cells xenografted into SCID-beige mice.

**Supplementary Fig. 3.**
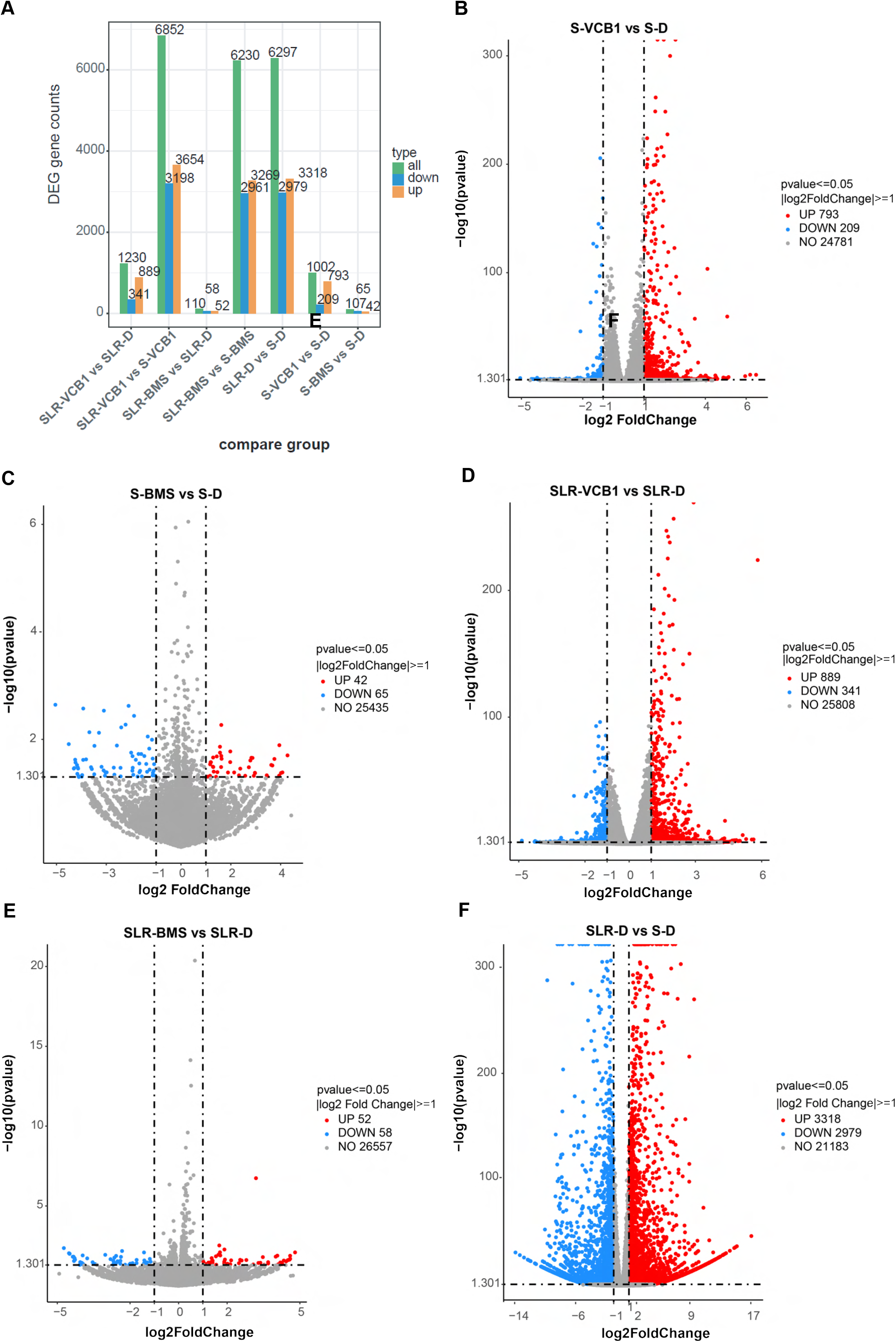
RNA sequencing identifies distinct transcriptional responses to VCB1 and BMS-303141 in ALK+ ALCL cells. (A) Summary of the total number and direction of differentially expressed genes identified across the indicated comparisons. (B) Comparison of the number of DEGs identified following VCB1 and BMS-303141 treatment in ALKi-sensitive SUPM2 parental cells. (C) Representative global transcriptional comparison of VCB1- and BMS-treated parental cells. (D) Differential gene expression analysis of S-VCB1 versus S-DMSO cells, identifying 793 upregulated and 209 downregulated genes using a threshold of log2 FC ≥1 and P<0.05. (E) Differential gene expression analysis of S-BMS versus S-DMSO cells, identifying 42 upregulated and 65 downregulated genes using the same criteria. (F) Differential gene expression analysis comparing untreated ALKi-resistant SUPM2-LR1000 (SLR) cells with parental SUPM2 cells, identifying 3,318 upregulated and 2,979 downregulated genes. Three biological replicates were analyzed per treatment group.

**Supplementary Fig. 4.**
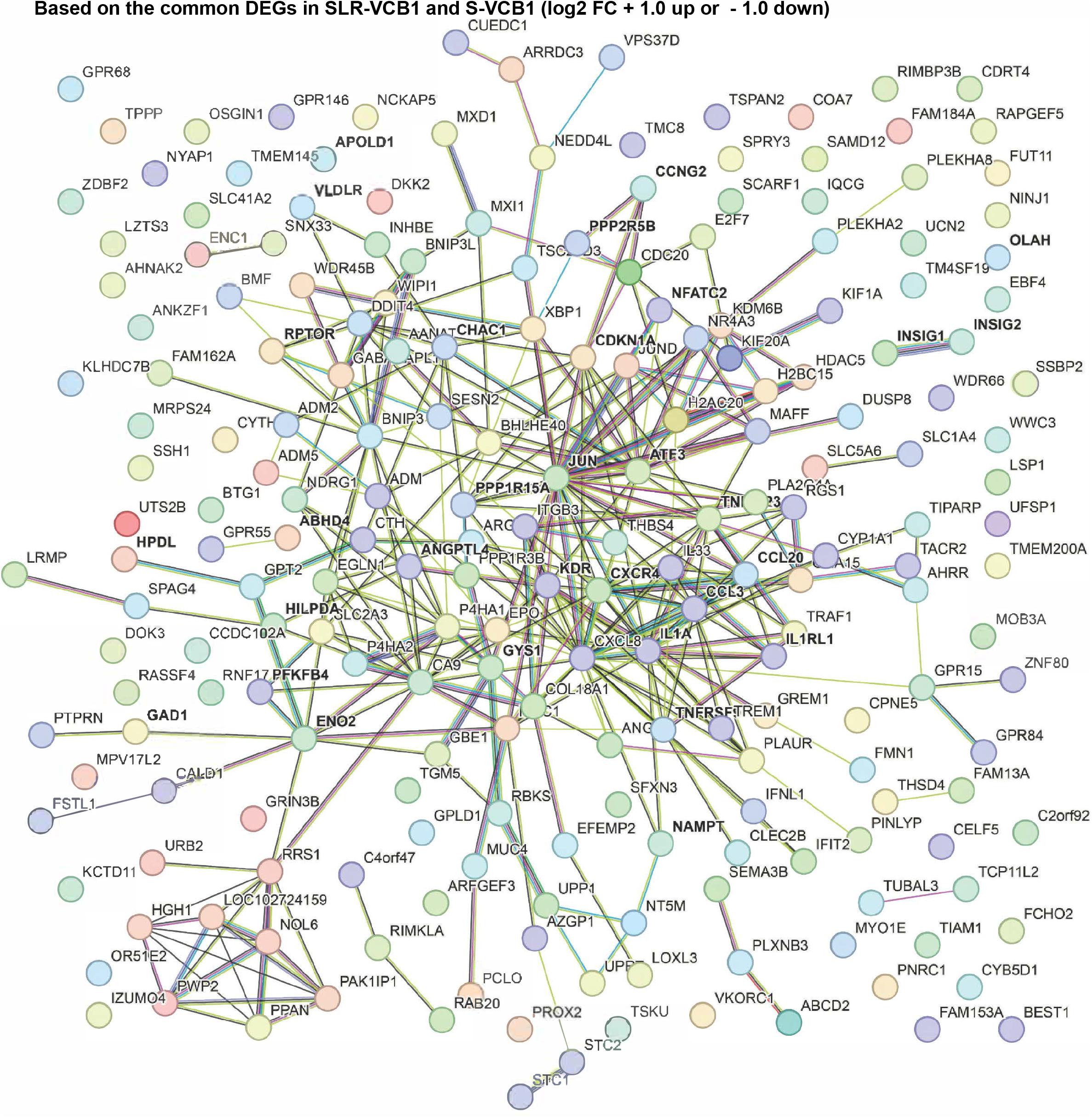
STRING-network analysis of common differentially expressed genes (DEGs) following VCB1 treatment. STRING-network analysis of DEGs commonly altered in TKI-sensitive and TKI-resistant cells using a log₂ fold-change (FC) threshold of ≥1.0 and P<0.05. The network illustrates functional interactions among common DEGs irrespective of biological function.

**Supplementary Fig. 5.**
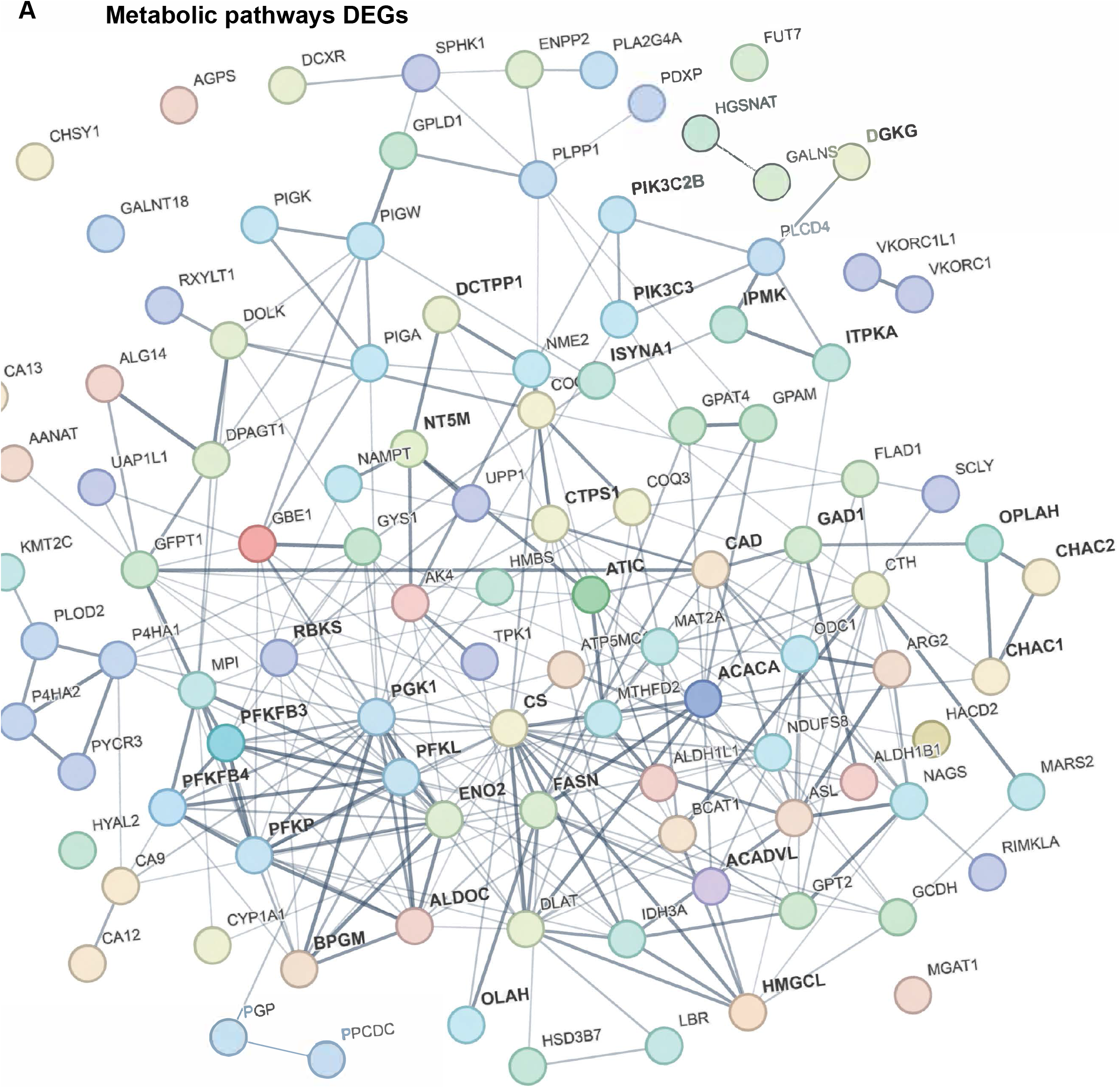
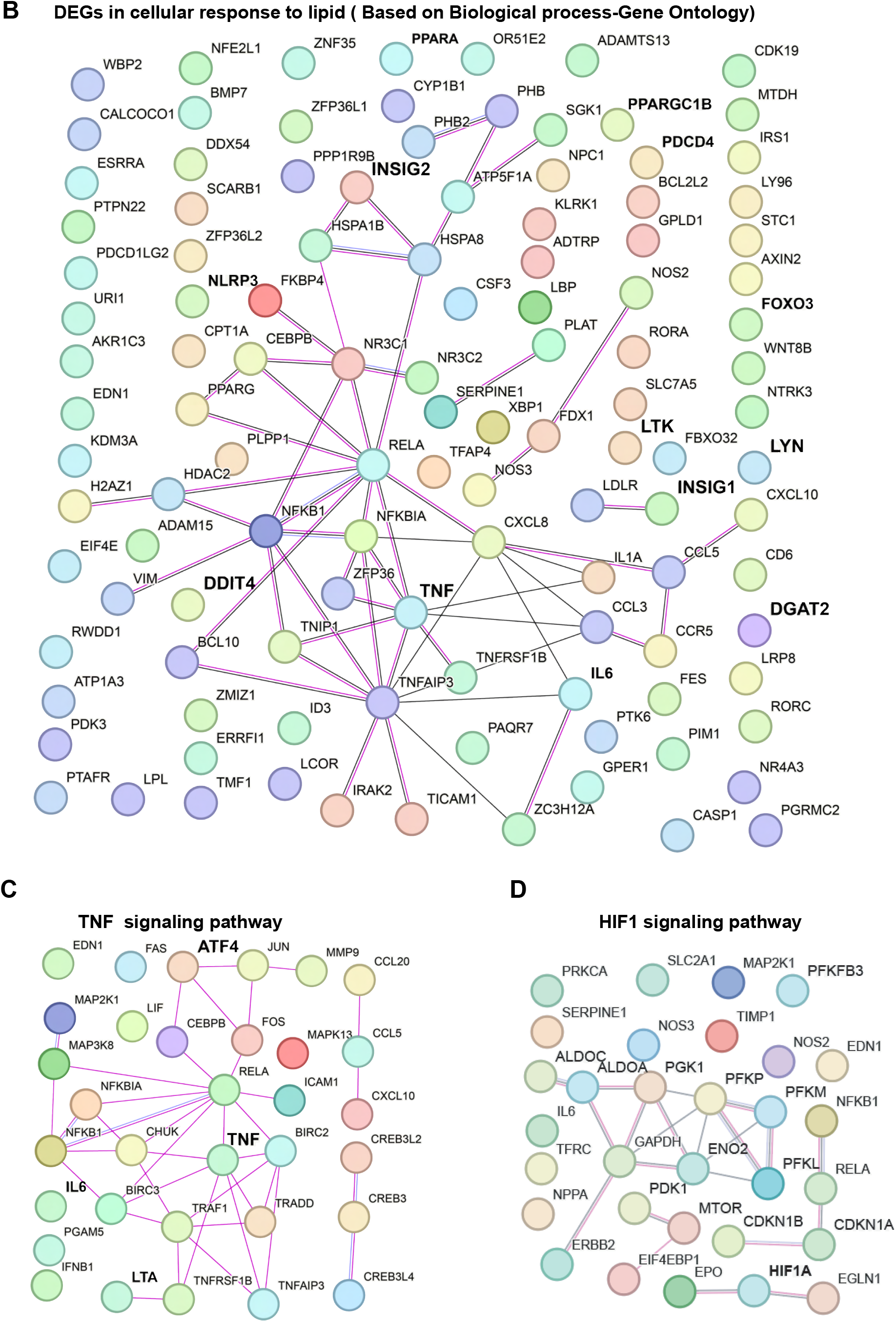
Functional network and pathway enrichment analysis of common VCB1-responsive DEGs. (A) STRING-network analysis of common DEGs associated with metabolic pathways using a log2 FC threshold of ≥1.0 and P<0.05. (B) Biological process enrichment of common DEGs associated with cellular lipid responses. (C, D) Network analysis showing TNF and HIF-1 signaling-associated transcriptional changes among common VCB1-responsive DEGs.

**Supplementary Fig. 6.**
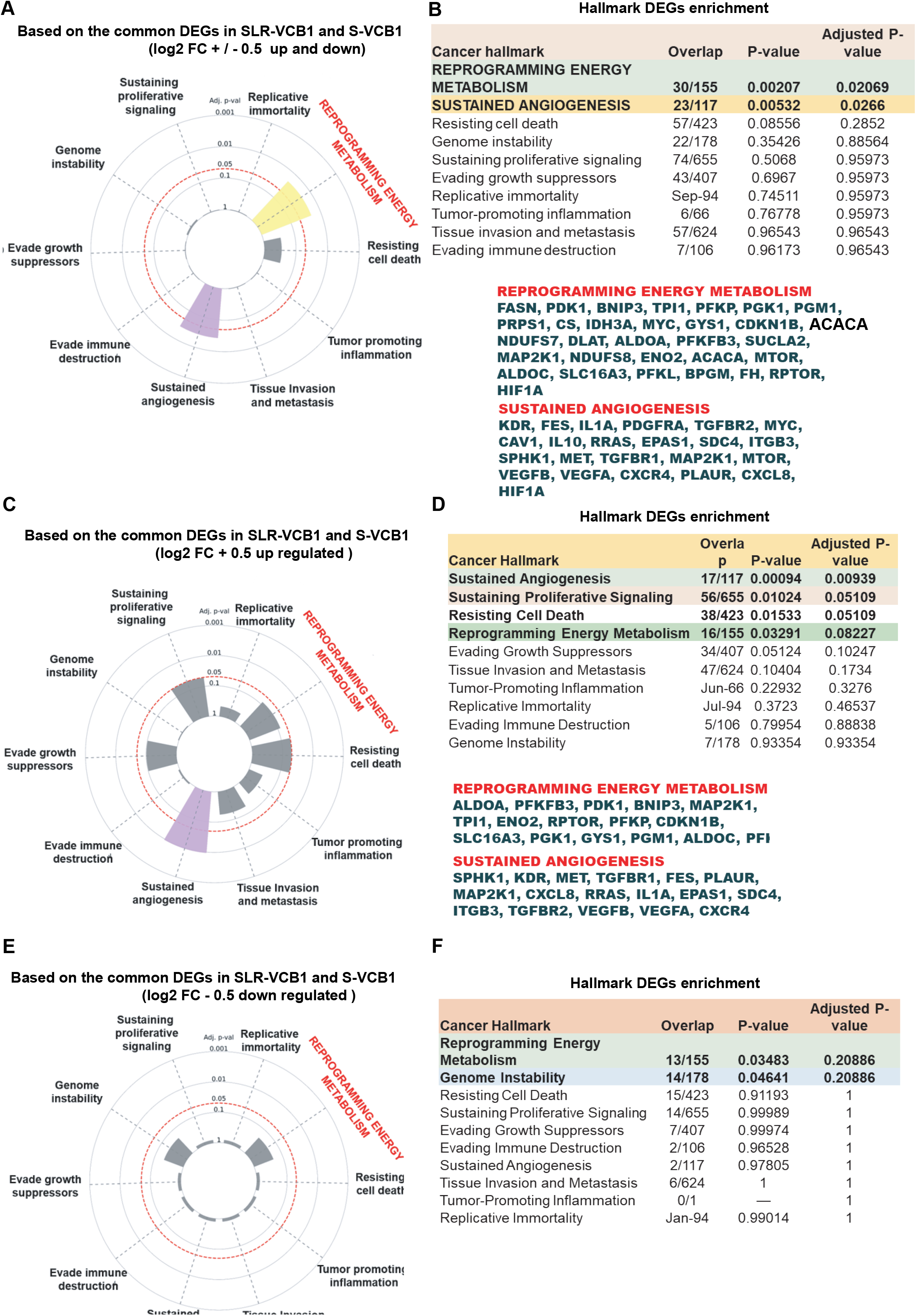
Hallmark cancer pathway enrichment among common VCB1-responsive DEGs. (A, B) Enrichment of reprogrammed energy metabolism and sustained angiogenesis among common DEGs identified using a log2 FC cutoff of ≥0.5. (C, D) Common upregulated DEGs were enriched for sustained angiogenesis, sustained proliferative signaling, resistance to cell death, and reprogrammed energy metabolism. (E, F) Common downregulated DEGs were predominantly enriched for reprogrammed energy metabolism.

**Supplementary Fig. 7.**
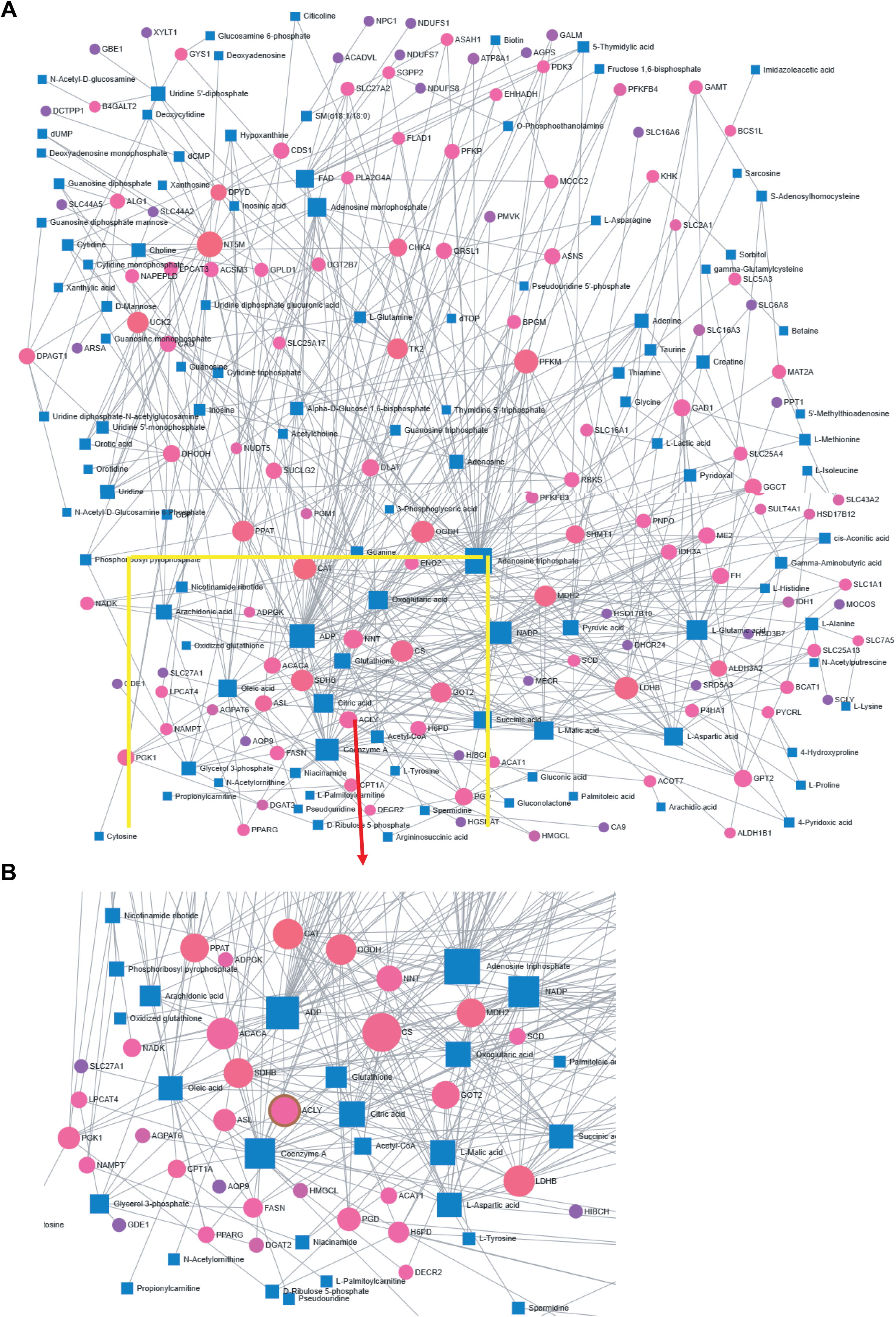
Gene-metabolite interaction network (A) Integrated gene-metabolite interaction network generated from VCB1-responsive transcriptomic and metabolomic datasets demonstrating enrichment of lipid-metabolic interactions in SLR cell line. (B) ACLY gene-metabolite specific network node highlighted from the above.

**Supplementary Table 1.**
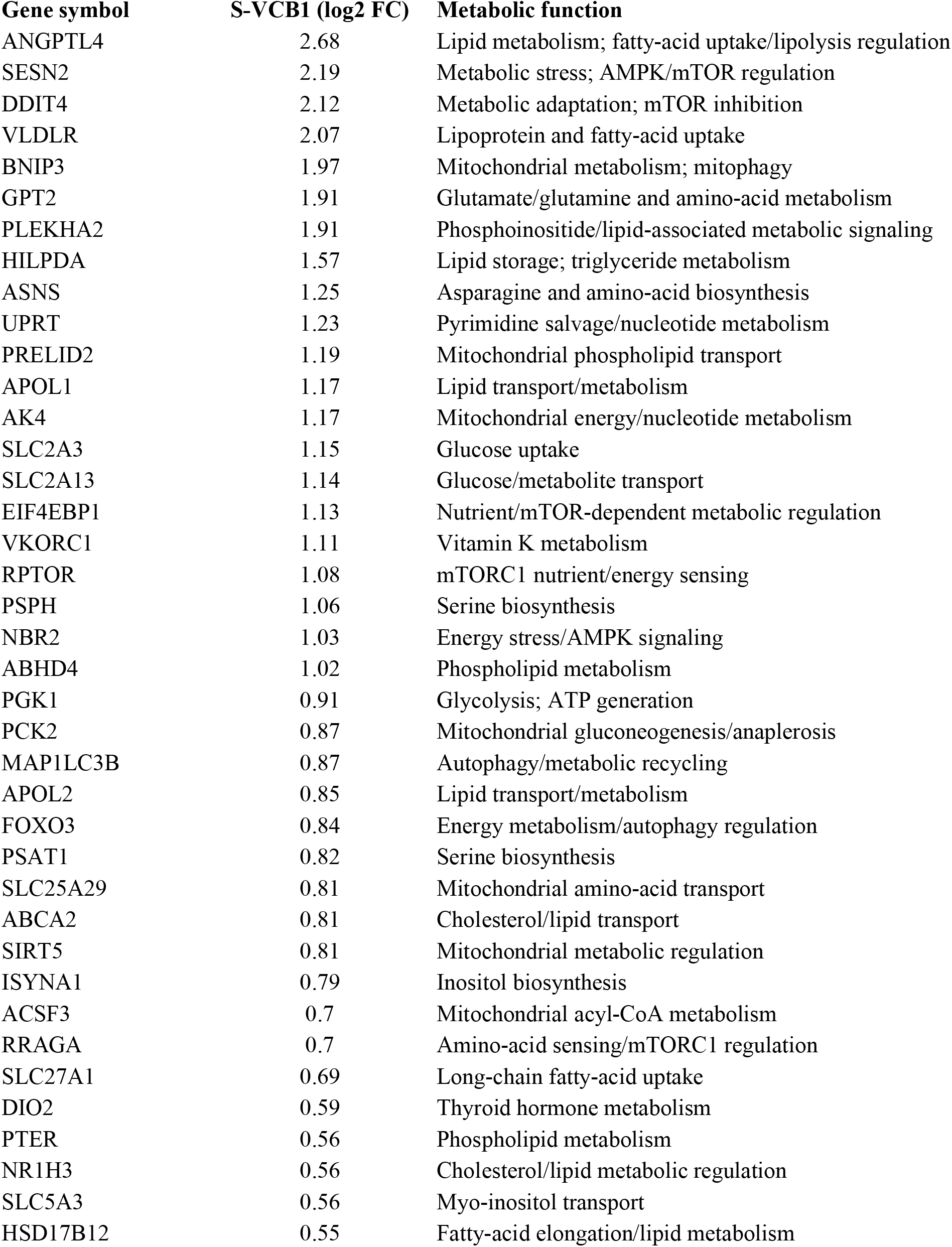
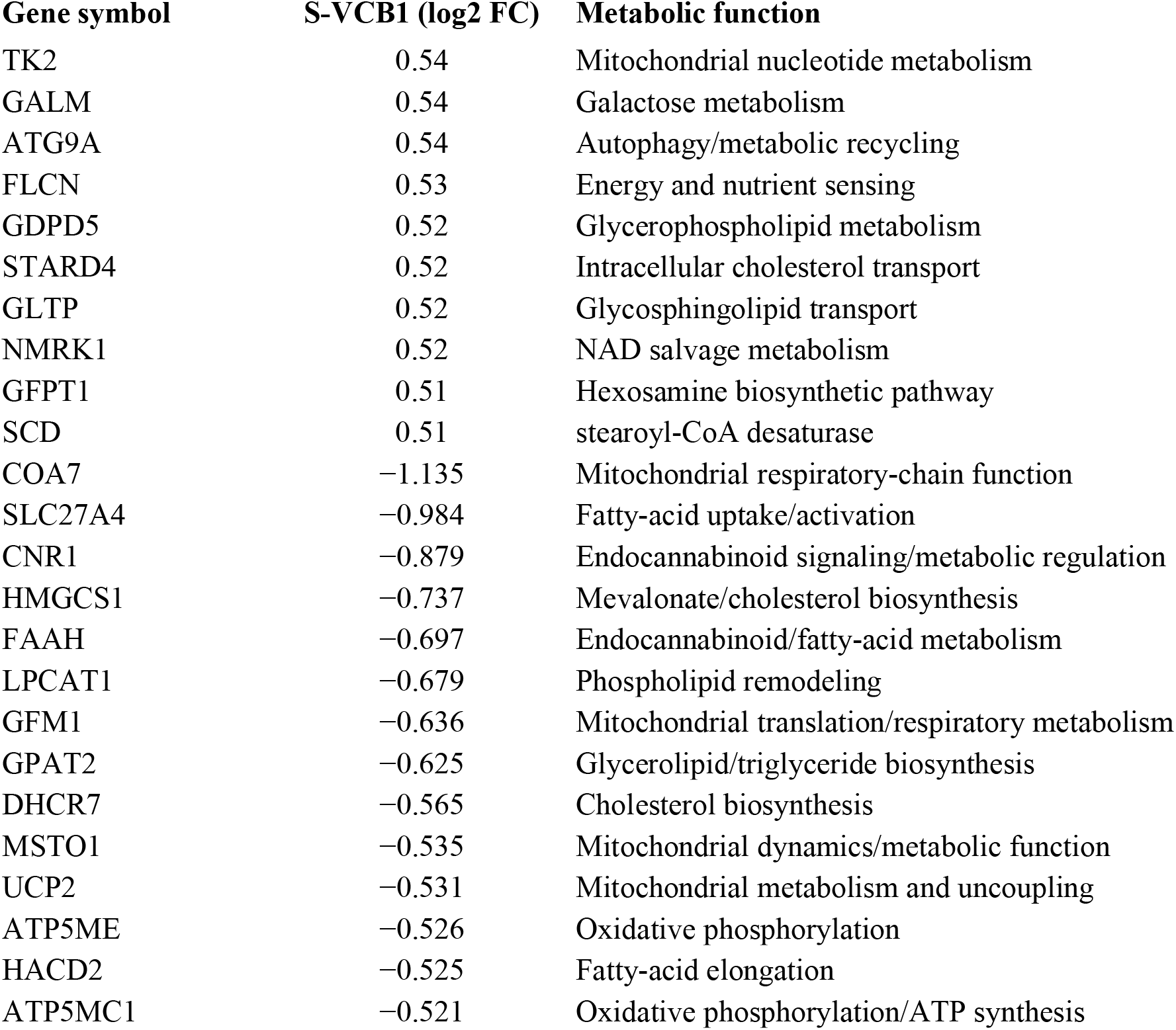
Metabolic function DEGs specific in S-VCB1 condition. Metabolic differentially expressed genes (DEGs) specific to SUPM2 parental cells following VCB1 treatment. SUPM2 parental-specific metabolic DEGs were identified using a log2 fold-change (FC) threshold of ≥ +0.5 or ≤ −0.5.

**Supplementary Table 2.** Metabolic function DEGs specific in SLR-VCB1 condition. Metabolic differentially expressed genes (DEGs) specific to SLR cells following VCB1 treatment. SLR cell-specific metabolic DEGs were identified using a log2 fold-change (FC) threshold of ≥ +0.5 or ≤ −0.5.

| **Gene symbol** | **SLR-VCB1 ( log2 FC)** | **Metabolic function** |
| --- | --- | --- |
| NT5M | -1.883 | Mitochondrial 5'-nucleotidase; nucleotide metabolism |
| SLC43A2 | -1.665 | Large neutral amino-acid transporter |
| EEF2KMT | -1.641 | Links protein synthesis to metabolic regulation |
| SULT4A1 | -1.622 | Sulfotransferase; small molecule/steroid metabolism |
| BCO2 | -1.379 | β-carotene/carotenoid metabolism |
| CHAC2 | -1.373 | Glutathione/cysteine metabolism |
| FXN | -1.348 | Fe-S cluster metabolism and oxidative phosphorylation |
| ATP8A1 | -1.327 | Phospholipid membrane metabolism |
| NDUFAF4 | -1.312 | Mitochondrial complex I assembly |
| SLC16A14 | -1.226 | Monocarboxylate/metabolite transport |
| CAD | -1.196 | De novo pyrimidine biosynthesis |
| DCTPP1 | -1.152 | dNTP metabolism |
| PGAM5 | -1.140 | Mitochondrial metabolism/stress signaling |
| SGPP2 | -1.127 | Sphingolipid metabolism |
| SLC16A9 | -1.127 | Monocarboxylate/carnitine-associated metabolite transport |
| FLAD1 | -1.122 | FAD/flavin metabolism |
| HSD17B8 | -0.910 | Steroid/lipid metabolism |
| CPT1A | -0.899 | Mitochondrial fatty-acid β-oxidation |
| CLUH | -0.885 | Mitochondrial protein/lipid metabolic homeostasis |
| NUDT5 | -0.881 | Nucleoside diphosphate/ADP-ribose metabolism |
| ALG1 | -0.879 | N-glycan/glycan metabolism |
| NT5DC3 | -0.877 | Nucleotide metabolism |
| ACSM3 | -0.859 | Fatty-acid activation to acyl-CoA |
| GOT2 | -0.857 | Amino-acid/TCA coupling |
| MTHFD1 | -0.848 | Folate-mediated one-carbon metabolism |
| LPCAT4 | -0.837 | Phospholipid remodeling |
| MAT2A | -0.826 | Methionine/SAM and methylation metabolism |
| UCK2 | -0.804 | Pyrimidine salvage |
| LPCAT3 | -0.803 | Phospholipid remodeling |
| PPAT | -0.795 | De novo purine biosynthesis |
| IDH3A | -0.779 | TCA cycle; α-ketoglutarate production |
| LDHB | -0.765 | Lactate ↔ pyruvate metabolism |
| CDS1 | -0.755 | CDP-diacylglycerol/phospholipid biosynthesis |
| OGDH | -0.749 | TCA cycle |
| PFKM | -0.748 | Glycolytic phosphofructokinase |
| ACAT1 | -0.739 | Acetyl-CoA, ketogenesis/cholesterol metabolism |
| FH | -0.734 | TCA cycle |
| NNT | -0.726 | Mitochondrial NADPH generation |
| MECR | -0.706 | Mitochondrial fatty-acid synthesis |
| PTDSS1 | -0.700 | Phosphatidylserine biosynthesis |
| MYLIP | -0.690 | Cholesterol homeostasis |
| NADK | -0.682 | NADP(H) production/redox metabolism |
| GDE1 | -0.680 | Glycerophospholipid/phosphoinositide metabolism |
| ALDH3A2 | -0.659 | Fatty-aldehyde oxidation |
| ASAH1 | -0.658 | Ceramide/sphingolipid metabolism |
| ABCD3 | -0.652 | Peroxisomal fatty-acid transport |
| SLC27A2 | -0.650 | Long-chain fatty-acid transport/activation |
| ARV1 | -0.641 | Sterol/lipid homeostasis |
| NDUFAF6 | -0.636 | Complex I assembly |
| HSD17B10 | -0.632 | Mitochondrial β-oxidation |
| NDUFAF8 | -0.630 | Complex I assembly |
| KHK | -0.629 | Fructose phosphorylation |
| EHHADH | -0.626 | Peroxisomal fatty-acid β-oxidation |
| GAMT | -0.626 | Creatine biosynthesis |
| MCAT | -0.625 | Mitochondrial fatty-acid synthesis |
| DECR2 | -0.624 | Peroxisomal fatty-acid β-oxidation |
| NDUFS7 | -0.624 | Mitochondrial complex I |
| SUCLG2 | -0.618 | TCA cycle/substrate-level phosphorylation |
| NAPEPLD | -0.616 | N-acyl-phosphatidylethanolamine/endocannabinoid metabolism |
| SHMT1 | -0.612 | Serine/one-carbon metabolism |
| SFXN4 | -0.608 | Mitochondrial metabolite/one-carbon transport |
| HACD3 | -0.604 | Fatty-acid elongation |
| CAT | -0.604 | H<sub>2</sub>O<sub>2</sub> detoxification/redox metabolism |
| ACOT7 | -0.602 | Acyl-CoA regulation |
| SDHB | -0.596 | TCA cycle/complex II |
| CHKA | -0.591 | Phosphatidylcholine synthesis |
| PMVK | -0.588 | Mevalonate pathway |
| FECH | -0.587 | Heme biosynthesis |
| DGAT2 | -0.582 | Triacylglycerol synthesis |
| DHCR24 | -0.579 | Cholesterol biosynthesis |
| IDH1 | -0.574 | Cytosolic isocitrate metabolism/NADPH |
| GPAM | -0.570 | Glycerolipid synthesis |
| NEU3 | -0.566 | Ganglioside/sialic-acid and lipid metabolism |
| MDH2 | -0.559 | TCA metabolism |
| SLC25A17 | -0.559 | Peroxisomal CoA/ATP transport |
| NDUFS1 | -0.557 | Mitochondrial complex I |
| PPT1 | -0.550 | Lysosomal lipid/protein depalmitoylation |
| HSD3B7 | -0.545 | Bile acid/cholesterol metabolism |
| HIBCH | -0.543 | Valine catabolism |
| MCCC2 | -0.543 | Branched-chain amino-acid metabolism |
| PGD | -0.541 | Pentose-phosphate pathway/NADPH |
| SLC25A13 | -0.540 | Mitochondrial aspartate/glutamate transport |
| ME2 | -0.537 | NADPH generation from malate |
| PDP2 | -0.534 | Pyruvate dehydrogenase activation |
| QRSL1 | -0.521 | Glutamine-dependent purine biosynthesis |
| MIOS | -0.520 | GATOR2/mTORC1 nutrient sensing |
| KLF16 | -0.520 | Metabolic/mitochondrial transcriptional regulation |
| STARD7 | -0.518 | Mitochondrial phospholipid transport |
| PAQR3 | -0.518 | Metabolic signaling/PI3K pathways |
| HTATIP2 | -0.518 | Mitochondrial/lipid metabolic stress regulation |
| RETSAT | -0.517 | Retinol metabolism |
| PNPO | -0.515 | Vitamin B6 metabolism |
| NIPA2 | -0.514 | Magnesium/metabolic regulation |
| PPARG | -0.513 | Lipid storage/utilization regulation |
| LDAH | -0.513 | Lipid droplet/lipid storage regulation |
| XYLT1 | -0.512 | Glycosaminoglycan biosynthesis |
| B4GALT2 | -0.511 | Glycan/galactose metabolism |
| DPAGT1 | -0.510 | N-glycosylation/carbohydrate metabolism |
| GGCT | -0.509 | Glutathione metabolism |
| DHODH | -0.505 | De novo pyrimidine synthesis |
| ACLY | -0.502 | Citrate → cytosolic acetyl-CoA; central carbon/lipid metabolism |
| SCAP | -0.502 | SREBP activation; cholesterol/lipid homeostasis |
| STOML2 | -0.502 | Mitochondrial membrane/metabolic signaling |
| DPYD | -0.500 | Pyrimidine catabolism |
| CHAC1 | 1.77 | Glutathione degradation/oxidative-stress response |
| GLUL | 1.5 | Glutamine synthesis/nitrogen metabolism |
| PDK1 | 0.99 | Pyruvate dehydrogenase inhibition/metabolic reprogramming |
| DMGDH | 0.93 | Glycine/dimethylglycine metabolism |
| SLC2A1 | 0.9 | Glucose uptake |
| SLC16A13 | 0.9 | Monocarboxylate/lipid metabolic regulation |
| AMPD3 | 0.9 | AMP/purine metabolism |
| STARD10 | 0.89 | Intracellular lipid handling/signaling |
| MSRB1 | 0.88 | Methionine/oxidative-redox metabolism |
| TCN2 | 0.85 | Vitamin B12 transport/cofactor homeostasis |
| DNAJB9 | 0.84 | ER proteostasis/metabolic stress |
| PFKFB3 | 0.82 | Glycolytic flux/glucose metabolism |
| GK | 0.81 | Glucose phosphorylation |
| SLC16A3 | 0.81 | Lactate export/glycolytic metabolism |
| AMPD2 | 0.8 | Purine nucleotide metabolism |
| SOD2 | 0.79 | Mitochondrial ROS detoxification |
| PDK3 | 0.76 | Suppresses pyruvate oxidation; glycolytic adaptation |
| CRELD2 | 0.76 | ER stress/metabolic adaptation |
| SLC25A36 | 0.74 | Mitochondrial nucleotide transport |
| ERO1A | 0.74 | ER redox metabolism |
| EDEM1 | 0.74 | ER-associated degradation |
| APOBR | 0.72 | Lipid uptake/metabolism |
| NAGK | 0.72 | Amino-sugar/amino-acid metabolism |
| OPLAH | 0.71 | Glutathione degradation |
| PGRMC2 | 0.69 | Lipid/cholesterol-associated metabolism |
| FUCA1 | 0.5256 | Glycan/carbohydrate degradation |
| GGCX | 0.5248 | Vitamin-K-dependent metabolic processing |
| C11orf54 | 0.5246 | pH/metabolic homeostasis |
| CLN8 | 0.5236 | ER-lysosome lipid trafficking |
| ARF4 | 0.5224 | Membrane/lipid trafficking |
| TG | 0.522 | Thyroid hormone precursor metabolism |
| IP6K1 | 0.5215 | Inositol phosphate/energy signaling |
| SLC24A1 | 0.521 | Ion/metabolic homeostasis |
| LYPLAL1 | 0.5209 | Lipid/lysophospholipid metabolism |
| CARS | 0.5205 | Cysteine/amino-acid utilization |
| AIFM2 | 0.5197 | Redox metabolism/lipid-peroxidation regulation |
| BLOC1S6 | 0.5193 | Endosomal/lysosomal trafficking; indirect lipid relevance |
| UGP2 | 0.5188 | UDP-glucose/carbohydrate metabolism |
| ADAM19 | 0.5178 | Extracellular protein processing; indirect metabolic relevance |
| GPR137 | 0.5172 | Lysosomal/nutrient-dependent metabolism |
| SMAP2 | 0.5165 | Membrane/lipid trafficking |
| SLC26A11 | 0.5113 | Anion/pH homeostasis affecting metabolism |
| CACNA1C | 0.5107 | Calcium signaling influencing mitochondrial metabolism |
| ENTPD7 | 0.51 | Extracellular nucleotide metabolism |
| PQLC1 | 0.5089 | Lysosomal amino-acid transport |
| OGT | 0.5088 | Nutrient-sensitive glucose/hexosamine signaling |
| BCAT1 | 0.5086 | Branched-chain amino-acid catabolism |
| PPM1K | 0.5084 | BCAA catabolism/BCKDH regulation |
| LACTB | 0.5069 | Mitochondrial/lipid metabolism |
| BCKDK | 0.5068 | BCAA oxidation regulation |
| HSPA13 | 0.5048 | Metabolic/proteostasis stress |
| OSBPL5 | 0.5037 | Oxysterol/lipid transport |
| TK1 | 0.5025 | Thymidine salvage/dTTP production |
| RAB3A | 0.5024 | Vesicular trafficking; indirect metabolic relevance |
| AK2 | 0.5006 | Mitochondrial ATP/ADP/AMP homeostasis |

## References

1. Wellen KE, Hatzivassiliou G, Sachdeva UM, Bui TV, Cross JR, Thompson CB. ATP-Citrate Lyase Links Cellular Metabolism to Histone Acetylation. Science. 2009;324(5930):1076–80.

2. Icard P, Wu Z, Fournel L, Coquerel A, Lincet H, Alifano M. ATP citrate lyase: A central metabolic enzyme in cancer. Cancer Lett. 2020;471:125–34.

3. Zaidi N, Swinnen JV, Smans K. ATP-citrate lyase: a key player in cancer metabolism. Cancer Res. 2012;72(15):3709–14.

4. Potapova IA, El-Maghrabi MR, Doronin SV, Benjamin WB. Phosphorylation of recombinant human ATP : citrate lyase by cAMP-dependent protein kinase abolishes homotropic allosteric regulation of the enzyme by citrate and increases the enzyme activity. Allosteric activation of ATP : citrate lyase by phosphorylated sugars. Biochemistry-Us. 2000;39(5):1169–79.

5. Basappa J, Citir M, Zhang Q, Wang HY, Liu XB, Melnikov O, et al. ACLY is the novel signaling target of PIP2/PIP3 and Lyn in acute myeloid leukemia. Heliyon. 2020;6(5).

6. Basappa J, ElAzzouny MA, Rolland DCM, Sahasrabuddhe AA, Ma K, Bazilevsky GA, et al. Tyrosine phosphorylation is critical for ACLY activity in lipid metabolism and cancer. bioRxiv. 2020:2020.01.20.910752.

7. Xie Z, Zhang M, Song Q, Cheng L, Zhang X, Song G, et al. Development of the novel ACLY inhibitor 326E as a promising treatment for hypercholesterolemia. Acta Pharm Sin B. 2023;13(2):739–53.

8. Morrow MR, Batchuluun B, Wu J, Ahmadi E, Leroux JM, Mohammadi-Shemirani P, et al. Inhibition of ATP-citrate lyase improves NASH, liver fibrosis, and dyslipidemia. Cell Metab. 2022;34(6):919–36 e8.

9. Convertini P, Santarsiero A, Todisco S, Gilio M, Palazzo D, Pappalardo I, et al. ACLY as a modulator of liver cell functions and its role in Metabolic Dysfunction-Associated Steatohepatitis. J Transl Med. 2023;21(1):568.

10. Migita T, Narita T, Nomura K, Miyagi E, Inazuka F, Matsuura M, et al. ATP Citrate Lyase: Activation and Therapeutic Implications in Non-Small Cell Lung Cancer. Cancer Research. 2008;68(20):8547–54.

11. Chen Y, Li K, Gong D, Zhang J, Li Q, Zhao G, et al. ACLY: A biomarker of recurrence in breast cancer. Pathol Res Pract. 2020;216(9):153076.

12. Shah S, Carriveau WJ, Li J, Campbell SL, Kopinski PK, Lim HW, et al. Targeting ACLY sensitizes castration-resistant prostate cancer cells to AR antagonism by impinging on an ACLY-AMPK-AR feedback mechanism. Oncotarget. 2016;7(28):43713–30.

13. Wang Y, Su K, Wang C, Deng T, Liu X, Sun S, et al. Chemotherapy-induced acetylation of ACLY by NAT10 promotes its nuclear accumulation and acetyl-CoA production to drive chemoresistance in hepatocellular carcinoma. Cell Death Dis. 2024;15(7):545.

14. Carrer A, Trefely S, Zhao S, Campbell SL, Norgard RJ, Schultz KC, et al. Acetyl-CoA Metabolism Supports Multistep Pancreatic Tumorigenesis. Cancer Discov. 2019;9(3):416–35.

15. Wen J, Min X, Shen M, Hua Q, Han Y, Zhao L, et al. ACLY facilitates colon cancer cell metastasis by CTNNB1. J Exp Clin Cancer Res. 2019;38(1):401.

16. Greenwood DL, Ramsey HE, Nguyen PTT, Patterson AR, Voss K, Bader JE, et al. Acly Deficiency Enhances Myelopoiesis through Acetyl Coenzyme A and Metabolic-Epigenetic Cross-Talk. Immunohorizons. 2022;6(12):837–50.

17. Andrieu GP, Hypolite G, Latiri M, Balducci E, Costa C, Verhoeyen E, et al. ATP citrate lyase is an essential player in the metabolic rewiring induced by PTEN loss during T-ALL development. Blood Adv. 2025;9(7):1670–91.

18. da Silva-Diz V, Singh A, Aleksandrova M, Kim O, Thai C, Lancho O, et al. A feedforward loop between ACLY and MYC supports T-ALL progression in vivo. Blood Neoplasia. 2025;2(2):100069.

19. Wang J, Ye W, Yan X, Guo Q, Ma Q, Lin F, et al. Low expression of ACLY associates with favorable prognosis in acute myeloid leukemia. J Transl Med. 2019;17(1):149.

20. Hatzivassiliou G, Zhao FP, Bauer DE, Andreadis C, Shaw AN, Dhanak D, et al. ATP citrate lyase inhibition can suppress tumor cell growth. Cancer Cell. 2005;8(4):311–21.

21. Hatzivassiliou G, Zhao F, Bauer DE, Andreadis C, Shaw AN, Dhanak D, et al. ATP citrate lyase inhibition can suppress tumor cell growth. Cancer Cell. 2005;8(4):311–21.

22. Pearce NJ, Yates JW, Berkhout TA, Jackson B, Tew D, Boyd H, et al. The role of ATP citrate-lyase in the metabolic regulation of plasma lipids. Hypolipidaemic effects of SB-204990, a lactone prodrug of the potent ATP citrate-lyase inhibitor SB-201076. Biochem J. 1998;334 ( Pt 1)(Pt 1):113–9.

23. Ma Z, Chu CH, Cheng D. A novel direct homogeneous assay for ATP citrate lyase. J Lipid Res. 2009;50(10):2131–5.

24. Wei J, Leit S, Kuai J, Therrien E, Rafi S, Harwood HJ, Jr., et al. An allosteric mechanism for potent inhibition of human ATP-citrate lyase. Nature. 2019;568(7753):566–70.

25. Liang JJ, Zhou XF, Long H, Li CY, Wei J, Yu XQ, et al. Recent advance of ATP citrate lyase inhibitors for the treatment of cancer and related diseases. Bioorg Chem. 2024;142:106933.

26. Thompson PD, Rubino J, Janik MJ, MacDougall DE, McBride SJ, Margulies JR, et al. Use of ETC-1002 to treat hypercholesterolemia in patients with statin intolerance. J Clin Lipidol. 2015;9(3):295-304.

27. Nissen SE, Lincoff AM, Brennan D, Ray KK, Mason D, Kastelein JJP, et al. Bempedoic Acid and Cardiovascular Outcomes in Statin-Intolerant Patients. N Engl J Med. 2023;388(15):1353–64.

28. Gautam J, Wu J, Lally JSV, McNicol JD, Fayyazi R, Ahmadi E, et al. ACLY inhibition promotes tumour immunity and suppresses liver cancer. Nature. 2025.

29. Verschueren KHG, Blanchet C, Felix J, Dansercoer A, De Vos D, Bloch Y, et al. Structure of ATP citrate lyase and the origin of citrate synthase in the Krebs cycle. Nature. 2019;568(7753):571–5.

30. Basappa J, ElAzzouny MA, Rolland DC, Velusamy T, Hwang S, Basrur V, et al. NPM-ALK Mediated Tyrosine Phosphorylation of ATP Citrate Lyase Regulates Lipid Metabolism and Promotes Oncogenesis of Anaplastic Large Cell Lymphoma. Blood. 2015;126(23):465-.

31. Pattabiraman N, Lobello C, Rushmore D, Nejati R, Mologni L, Wasik M, et al. Structure-guided discovery of potent acly inhibitors with therapeutic potential in ALK plus anaplastic large cell lymphoma and other hematologic malignancies. Blood. 2025;146:6815-.

32. Basappa J, Citir M, Zhang Q, Wang HY, Liu X, Melnikov O, et al. ACLY is the novel signaling target of PIP(2)/PIP(3) and Lyn in acute myeloid leukemia. Heliyon. 2020;6(5):e03910.

33. Ceccon M, Mologni L, Bisson W, Scapozza L, Gambacorti-Passerini C. Crizotinib-resistant NPM-ALK mutants confer differential sensitivity to unrelated Alk inhibitors. Mol Cancer Res. 2013;11(2):122–32.

34. Pattabiraman N, Lobello C, Rushmore D, Mologni L, Wasik M, Basappa J. A Novel Allosteric Inhibitor Targeting IMPDH at Y233 Overcomes Resistance to Tyrosine Kinase Inhibitors in Lymphoma. Cancers (Basel). 2025;17(20).

35. Basappa J, Goldman AR, Lobello C, Wang S, Rushmore D, Melnikov O, et al. Mitochondrial ACSS1 links acetate metabolism to pyrimidine biosynthesis in nutrient-stressed B-cell lymphomas. Cancer Lett. 2026;649:218488.

36. Di Marcantonio D, Martinez E, Kanefsky JS, Huhn JM, Gabbasov R, Gupta A, et al. ATF3 coordinates serine and nucleotide metabolism to drive cell cycle progression in acute myeloid leukemia. Mol Cell. 2021;81(13):2752–64 e6.

37. Li J, Agarwal E, Bertolini I, Seo JH, Caino MC, Ghosh JC, et al. The mitophagy effector FUNDC1 controls mitochondrial reprogramming and cellular plasticity in cancer cells. Sci Signal. 2020;13(642).

38. Lombardi S, Goldman AR, Tang HY, Kossenkov AV, Liu H, Zhou W, et al. Targeting Fatty Acid Reprogramming Suppresses CARM1-expressing Ovarian Cancer. Cancer Res Commun. 2023;3(6):1067–77.

39. Zaidi N, Royaux I, Swinnen JV, Smans K. ATP citrate lyase knockdown induces growth arrest and apoptosis through different cell- and environment-dependent mechanisms. Mol Cancer Ther. 2012;11(9):1925–35.

40. da Silva-Diz V, Singh A, Aleksandrova M, Kim O, Thai C, Lancho O, et al. Therapeutic targeting of ACLY in T-ALL in vivo. bioRxiv. 2024.

41. Huang da W, Sherman BT, Zheng X, Yang J, Imamichi T, Stephens R, et al. Extracting biological meaning from large gene lists with DAVID. Curr Protoc Bioinformatics. 2009;Chapter 13:Unit 13 1.

42. Wellen KE, Hatzivassiliou G, Sachdeva UM, Bui TV, Cross JR, Thompson CB. ATP-citrate lyase links cellular metabolism to histone acetylation. Science. 2009;324(5930):1076–80.

43. Zhao S, Torres A, Henry RA, Trefely S, Wallace M, Lee JV, et al. ATP-Citrate Lyase Controls a Glucose-to-Acetate Metabolic Switch. Cell Rep. 2016;17(4):1037–52.

44. Chen Q, Du J, Cui K, Fang W, Zhao Z, Chen Q, et al. Acetyl-CoA derived from hepatic mitochondrial fatty acid beta-oxidation aggravates inflammation by enhancing p65 acetylation. iScience. 2021;24(11):103244.

45. Kaymak I, Watson MJ, Oswald BM, Ma S, Johnson BK, DeCamp LM, et al. ACLY and ACSS2 link nutrient-dependent chromatin accessibility to CD8 T cell effector responses. J Exp Med. 2024;221(9).

46. Pang Z, Lu Y, Zhou G, Hui F, Xu L, Viau C, et al. MetaboAnalyst 6.0: towards a unified platform for metabolomics data processing, analysis and interpretation. Nucleic Acids Res. 2024;52(W1):W398–W406.

47. Ben-Sahra I, Hoxhaj G, Ricoult SJH, Asara JM, Manning BD. mTORC1 induces purine synthesis through control of the mitochondrial tetrahydrofolate cycle. Science. 2016;351(6274):728–33.

48. Donnelly N, Gorman AM, Gupta S, Samali A. The eIF2alpha kinases: their structures and functions. Cell Mol Life Sci. 2013;70(19):3493–511.

49. Labbe K, LeBon L, King B, Vu N, Stoops EH, Ly N, et al. Specific activation of the integrated stress response uncovers regulation of central carbon metabolism and lipid droplet biogenesis. Nat Commun. 2024;15(1):8301.

50. Ryoo HD. The integrated stress response in metabolic adaptation. J Biol Chem. 2024;300(4):107151.

51. Thomas-Jardin S, Suresh S, Arce A, Novaresi N, Deng Q, Stein E, et al. The Integrated Stress Response Pathway Coordinates Translational Control of Multiple Immune Checkpoints in Lung Cancer. Cancer Res. 2025;85(14):2574–90.

52. Lu Q, Tang S, Fang S, Wu Y, Chen L, Liang M, et al. Multi-omics profiling of the diabetic human heart reveals coupled dysregulation in lipid metabolism, mitophagy, and extracellular matrix remodeling. Genome Med. 2026;18(1).

53. Zheng Y, Zhou Q, Zhao C, Li J, Yu Z, Zhu Q. ATP citrate lyase inhibitor triggers endoplasmic reticulum stress to induce hepatocellular carcinoma cell apoptosis via p-eIF2alpha/ATF4/CHOP axis. J Cell Mol Med. 2021;25(3):1468–79.

54. Oliveira MM, Mohamed M, Elder MK, Banegas-Morales K, Mamcarz M, Lu EH, et al. The integrated stress response effector GADD34 is repurposed by neurons to promote stimulus-induced translation. Cell Rep. 2024;43(2):113670.

55. Sun J, Ren H, Wang J, Xiao X, Zhu L, Wang Y, et al. CHAC1: a master regulator of oxidative stress and ferroptosis in human diseases and cancers. Front Cell Dev Biol. 2024;12:1458716.

56. Xiang W, Lv H, Xing F, Sun X, Ma Y, Wu L, et al. Inhibition of ACLY overcomes cancer immunotherapy resistance via polyunsaturated fatty acids peroxidation and cGAS-STING activation. Sci Adv. 2023;9(49):eadi2465.

57. Jia H, Ma L, Liu J, Gao M, Liang X, Zhang F, et al. Organelle stress in NLRP3 inflammasome: a central mediator of neurodegenerative diseases. Mol Neurodegener. 2026;21(1):8.

58. Ray KK, Bays HE, Catapano AL, Lalwani ND, Bloedon LT, Sterling LR, et al. Safety and Efficacy of Bempedoic Acid to Reduce LDL Cholesterol. N Engl J Med. 2019;380(11):1022–32.

59. Pinkosky SL, Newton RS, Day EA, Ford RJ, Lhotak S, Austin RC, et al. Liver-specific ATP-citrate lyase inhibition by bempedoic acid decreases LDL-C and attenuates atherosclerosis. Nat Commun. 2016;7:13457.

60. Gautam J, Wu J, Lally JSV, McNicol JD, Fayyazi R, Ahmadi E, et al. ACLY inhibition promotes tumour immunity and suppresses liver cancer. Nature. 2025;645(8080):507–17.

61. Di Pastena F, Gautam J, Lally JSV, Fayyazi R, Grasset E, Bhattacharya D, et al. Dual inhibition of ACLY and ACSS2 by EVT0185 reduces steatosis, hepatic stellate cell activation, and fibrosis in mouse models of MASH. Cell Metab. 2026;38(1):33–49 e10.

62. Piktel D, Nair RR, Rellick SL, Geldenhuys WJ, Martin KH, Craig MD, et al. Pitavastatin Is Anti-Leukemic in a Bone Marrow Microenvironment Model of B-Lineage Acute Lymphoblastic Leukemia. Cancers (Basel). 2022;14(11).

63. Wartewig T, Daniels J, Schulz M, Hameister E, Joshi A, Park J, et al. PD-1 instructs a tumor-suppressive metabolic program that restricts glycolysis and restrains AP-1 activity in T cell lymphoma. Nat Cancer. 2023;4(10):1508–25.

64. Petovari G, Danko T, Tokes AM, Vetlenyi E, Krencz I, Raffay R, et al. In Situ Metabolic Characterisation of Breast Cancer and Its Potential Impact on Therapy. Cancers (Basel). 2020;12(9).

65. Hart ML, Quon E, Vigil ABG, Engstrom IA, Newsom OJ, Davidsen K, et al. Mitochondrial redox adaptations enable alternative aspartate synthesis in SDH-deficient cells. Elife. 2023;12.

66. Sola-Garcia A, Caliz-Molina MA, Espadas I, Petr M, Panadero-Moron C, Gonzalez-Moran D, et al. Metabolic reprogramming by Acly inhibition using SB-204990 alters glucoregulation and modulates molecular mechanisms associated with aging. Commun Biol. 2023;6(1):250.

67. Granchi C. ATP citrate lyase (ACLY) inhibitors: An anti-cancer strategy at the crossroads of glucose and lipid metabolism. Eur J Med Chem. 2018;157:1276–91.

68. Chambers KT, Chen Z, Lai L, Leone TC, Towle HC, Kralli A, et al. PGC-1beta and ChREBP partner to cooperatively regulate hepatic lipogenesis in a glucose concentration-dependent manner. Mol Metab. 2013;2(3):194–204.

69. Zhang H, Li L, Chen Q, Li M, Feng J, Sun Y, et al. PGC1beta regulates multiple myeloma tumor growth through LDHA-mediated glycolytic metabolism. Mol Oncol. 2018;12(9):1579–95.

70. Korobkina ED, Calejman CM, Haley JA, Kelly ME, Li H, Gaughan M, et al. Brown fat ATP-citrate lyase links carbohydrate availability to thermogenesis and guards against metabolic stress. Nat Metab. 2024;6(11):2187–202.

71. Huhn JM, Valin L, Basse M, Sokei J, Gabriel N, Martinez E, et al. ATF4 coordinates amino acid and nucleotide synthesis with selective protein translation to ensure proper DNA replication timing in leukemia cells. Nat Commun. 2026;17(1).

72. Watanabe A, Tipgomut C, Totani H, Yoshimura K, Iwano T, Bashiri H, et al. Noncanonical TCA cycle fosters canonical TCA cycle and mitochondrial integrity in acute myeloid leukemia. Cancer Sci. 2025;116(1):152–63.

73. Li Z, Deng W, Yang L, Tang C, Yue JM, Monteiro O, et al. Lipid metabolism in homeostasis and disease. Signal Transduct Target Ther. 2026;11(1):55.

74. Wang Q, Yang HS. The role of Pdcd4 in tumour suppression and protein translation. Biol Cell. 2018.

75. Matsuhashi S, Manirujjaman M, Hamajima H, Ozaki I. Control Mechanisms of the Tumor Suppressor PDCD4: Expression and Functions. Int J Mol Sci. 2019;20(9).

76. Hilliard A, Hilliard B, Zheng SJ, Sun H, Miwa T, Song W, et al. Translational regulation of autoimmune inflammation and lymphoma genesis by programmed cell death 4. J Immunol. 2006;177(11):8095–102.

